# A Novel SARS-CoV-2 Multitope Protein/Peptide Vaccine Candidate is Highly Immunogenic and Prevents Lung Infection in an AAV hACE2 Mouse Model and non-human primates

**DOI:** 10.1101/2020.11.30.399154

**Authors:** Farshad Guirakhoo, Lucy Kuo, James Peng, Juin-Hua Huang, Be-Shen Kuo, Feng Lin, Yaw-Jen Liu, Zhi Liu, Grace Wu, Shuang Ding, Kou-Liang Hou, Jennifer Cheng, Vicky Yang, Hank Jiang, Jason Wang, Tony Chen, WeiGuo Xia, Ed Lin, Chung Ho Hung, Hui-Jung Chen, Zhonghao Shih, Yi-Ling Lin, Shixia Wang, Valorie Ryan, Brandon T. Schurter, Mei Mei Hu, Gray Heppner, Delphine C. Malherbe, Alexander Bukreyev, Michael Hellerstein, Thomas P. Monath, Chang Yi Wang

## Abstract

A novel multitope protein-peptide vaccine against Severe Acute Respiratory Syndrome Coronavirus 2 (SARS-CoV-2) infection and disease is described in this report. The initial development and characterization experiments are presented along with proof-of-concept studies for the vaccine candidate UB-612. UB-612 consists of eight components rationally designed for induction of potently neutralizing antibodies and broad T cell responses against SARS-CoV-2: the S1-RBD-sFc fusion protein, six synthetic peptides (one universal peptide and five SARS-CoV-2-derived peptides), a proprietary CpG TLR-9 agonist at low concentration as an excipient, and aluminum phosphate adjuvant. Through immunogenicity studies in Guinea pigs and rats, we optimized the design of protein/peptide immunogens and selected an adjuvant system, yielding a vaccine that provides excellent S1-RBD binding and high neutralizing antibody responses, robust cellular responses, and a Th1-oriented response at low doses. In challenge studies, UB- 612 vaccination reduced viral load and prevented development of disease in mouse and non-human primate challenge models. With a Phase 1 trial completed, a Phase 2 trial ongoing in Taiwan, and additional trials planned to support global authorizations, UB-612 is a highly promising and differentiated vaccine candidate for prevention of SARS-CoV-2 infection and COVID-19 disease.

**Author Summary:** SARS-CoV-2 virus, the causative agent of Coronavirus Disease 2019 (COVID-19), has spread globally since its origin in 2019, causing an unprecedented public health crisis that has resulted in greater than 4.7 million deaths worldwide. Many vaccines are under development to limit disease spread and reduce the number of cases, but additional candidates that promote a robust immune response are needed. Here, we describe a multitope protein-peptide vaccine platform that is unique among COVID-19 vaccines. The advantages of our approach are induction of both high levels of neutralizing antibodies as well as a Th/CTL response in the vaccinated host, which mimics the immune response that occurs after natural infection with SARS-CoV-2. We demonstrate that our vaccine is immunogenic and effective in preventing disease in several animal models, including AAV-hACE-2 transduced mice, and both rhesus and cynomolgus macaques. Importantly, no immunopathology was observed in the lungs of immunized animals, therefore showing that antibody-dependent enhancement (ADE) does not occur. Our study provides an additional, novel vaccine candidate for advancement in clinical trials to treat and prevent SARS-CoV-2 infection and COVID-19 disease.

## Introduction

A novel coronavirus, Severe Acute Respiratory Syndrome Coronavirus 2 (SARS-CoV-2) was identified as the causative agent of a cluster of cases of the new coronavirus disease 2019 (COVID-19), in Wuhan China, in December 2019 [1]. SARS-CoV-2 has caused over 229 million cases of COVID-19 and greater than 4.7 million deaths worldwide as of September 21, 2021 [2]. In response to this unprecedented public health crisis, many vaccine platforms are under development, including inactivated virus, recombinant adenovirus-based vectors, recombinant proteins, and nucleic acid approaches. This paper describes a novel multitope protein-peptide vaccine candidate. The UB-612 vaccine is unique in that it has been developed specifically to address the need for a vaccine that elicits a strong, neutralizing antibody response targeting the receptor binding domain (RBD) of the Spike protein while simultaneously stimulating T cell responses to conserved peptides derived from three structural proteins: S2 subunit of Spike, Membrane (M) and Nucleocapsid (N) of the virus. UB-612 elicits high levels of neutralizing antibodies in addition to a Th1 prone immune response. Unlike some other full-length S protein- based vaccines, the RBD-focused antibody responses induced for UB-612 vaccine can avoid any potential antibody-dependent enhancement (ADE) in the lungs [3, 4].

The S1-RBD is a critical component of SARS-CoV-2, as it is required for cell attachment and represents the principal neutralizing domain of the virus [5–7]. Use of this truncated portion of the S protein as the vaccine antigen could provide a margin of safety not achievable with the full-length S protein and thereby eliminating the possibility of potentially deadly side effects that have previously been shown in animal models with SARS-CoV and MERS as well as with an inactivated RSV vaccine resulting in its withdrawal from the market [4,8–11]. Therefore, we chose to investigate S1-RBD as a potential immunogen for our novel UB-612 vaccine, and ultimately identified S1-RBD-sFc as the primary immunogen for induction of neutralizing antibodies and a memory B cell response. S1-RBD-sFc is a recombinant protein made through fusion of S1-RBD of SARS-CoV-2 to a single chain fragment crystallizable region (sFc) of a human IgG1. Genetic fusion of a vaccine antigen to a Fc fragment has been successfully shown to promote antibody induction and neutralizing activity, for example against HIV gp120 in Rhesus macaques or Epstein Barr virus gp350 in BALB/c mice [12–13]. Moreover, engineered Fc has been used in many therapeutic antibodies as a solution to minimize non-specific binding, increase solubility, yield, thermostability, and *in vivo* half-life [14]. Furthermore, the Fc-tagged RBD allows purification with Protein A, Protein G or Protein L affinity columns, yielding a high quality and low-cost purified product.

The durability of the antibody response after SARS-CoV-2 infection is unknown, with several studies demonstrating variable lengths of persistence for neutralizing antibody titers. One study found that the IgG response to S protein declined rapidly in >90% of SARS-CoV-2 infected individuals within 2-3 months [15, 16]. Additionally, a neutralizing response against the S protein alone is unlikely to provide lasting protection against SARS-CoV-2 and its emerging variants with mutated B-cell epitopes [17]. However, other studies have shown relatively stable antibody titers for up to 3-4 months after SARS-CoV-2 infection [18, 19]. Memory T cells to SARS-CoV-1 were also shown to persist 11-17 years after the original SARS outbreak in 2003 [20, 21]. Because the vast majority of reported CD8+ T cell epitopes in SARS-CoV-2 proteins are located in the ORF1ab, N, M, and ORF3a regions [22], while only three are present in the S protein, we included Th/CTL epitopes from highly conserved sequences derived from all three proteins (S, nucleocapsid, N and membrane, M) of SARS-CoV-2 [23–30] in the design of our UB-612 vaccine.

To enhance the immune response of the antigen portion (S1-RBD-sFc), we added our proprietary peptide UBITh®1a to the Th/CTL peptide mixture. UBITh®1a is a proprietary synthetic peptide with an original framework sequence derived from the measles virus fusion protein (MVF) modified to allow accommodation of multiple MHC class II binding motifs. In previous studies, attachment of UBITh®1a to a target “functional B epitope peptide” derived from a self-protein rendered the self-peptide immunogenic, thus breaking immune tolerance [31]. Proprietary CpG oligonucleotide (CpG1) [32] at low concentration is included as excipient to bring the rationally designed immunogens together through “charge neutralization” to stabilize the Th and CTL peptides by dipolar interactions between the negatively charged CpG1 molecule and positively charged peptides. In addition, activation of TLR-9 signaling by CpG is known to promote IgA production and favor the Th1 immune response [33]. The UBITh®1a peptide is incorporated as one of the Th peptides for its “epitope cluster” nature to further enhance the antiviral activity of the SARS-CoV-2 derived Th and CTL epitope peptides UB-612 includes, in addition to the recombinant S1-RBD-sFc fusion protein, CTL and Th epitope peptides selected from immunodominant S2, M and N regions known to bind to human MHC I and II molecules. Most reactivity to Spike protein comes from CD4+ T cells, and there is only one reported dominant CD8+ T cell epitope in the S protein, which resides within the RBD [22]. The smaller M and N structural proteins are recognized by T cells of patients who successfully controlled their infection [22, 29]. The five Th and CTL epitope peptides are selected from sequences of S2, M and N proteins of SARS-CoV-2, while the UBITh^®^1a is a proprietary T helper peptide adapted from measles virus fusion (MVF) protein. This results in balanced B cells (induction of neutralizing antibodies) and Th/CTL responses in a vaccinated host. This mixture of S1-RBD-sFc and Th/CTL peptides is designed to elicit T cell activation, memory B cell recall and effector functions similar to those elicited after natural SARS-CoV-2 infection.

Finally, to further improve the immune response, UB-612 is formulated with an aluminum phosphate (Adju-Phos®) adjuvant, which promotes Th2 responses via the nucleotide binding oligomerization domain (NOD) like receptor protein 3 (NLRP3) inflammasome pathway. Additionally, it has pro-phagocytic and repository effects with a long record of safety and the ability to improve immune responses to target proteins in many vaccine formulations [34, 35].

In this paper, we describe selection of the S1-RBD-sFc protein from among three candidates with different Fc-fusion structures. Guinea pigs (GP) were vaccinated with one of the three constructs transiently expressed in CHO cells. The lead candidate was chosen, based on highest neutralization and S1:ACE2:binding inhibition titers, and was further formulated with Th/CTL peptides (UB-612) [36]. We then confirmed immunogenicity and efficacy in AAV-hACE-2 transduced mice, as well as rhesus and cynomolgus macaques.

## Results

### Construction and characterization of S1-RBD-sFc

The UB-612 vaccine immunogen was designed to contain an S1-RBD-sFc fusion protein plus five synthetic Th/CTL peptides for class I and II MHC molecules derived from SARS-CoV-2 S2, M, and N proteins. To identify the best RBD immunogen to induce neutralizing antibody responses, three S1-RBD-based protein antigen (sequences aa340-539) vaccine candidates were designed: S1-RBD-sFc (single chain Fc), S1-RBDa-sFc (RBD domain modified to reduce a Cys-disulfide bond for better domain folding), and S1- RBD-Fc (double chain Fc) (structure of S1-RBD-sFc illustrated in **Fig 1B**). These synthetic genes were transfected into Chinese Hamster Ovary (CHO) cells for transient expression of proteins for initial studies. The immunogenicity of each vaccine candidate was tested in GPs, to select the lead B cell immunogen candidate.

**Fig 1.**
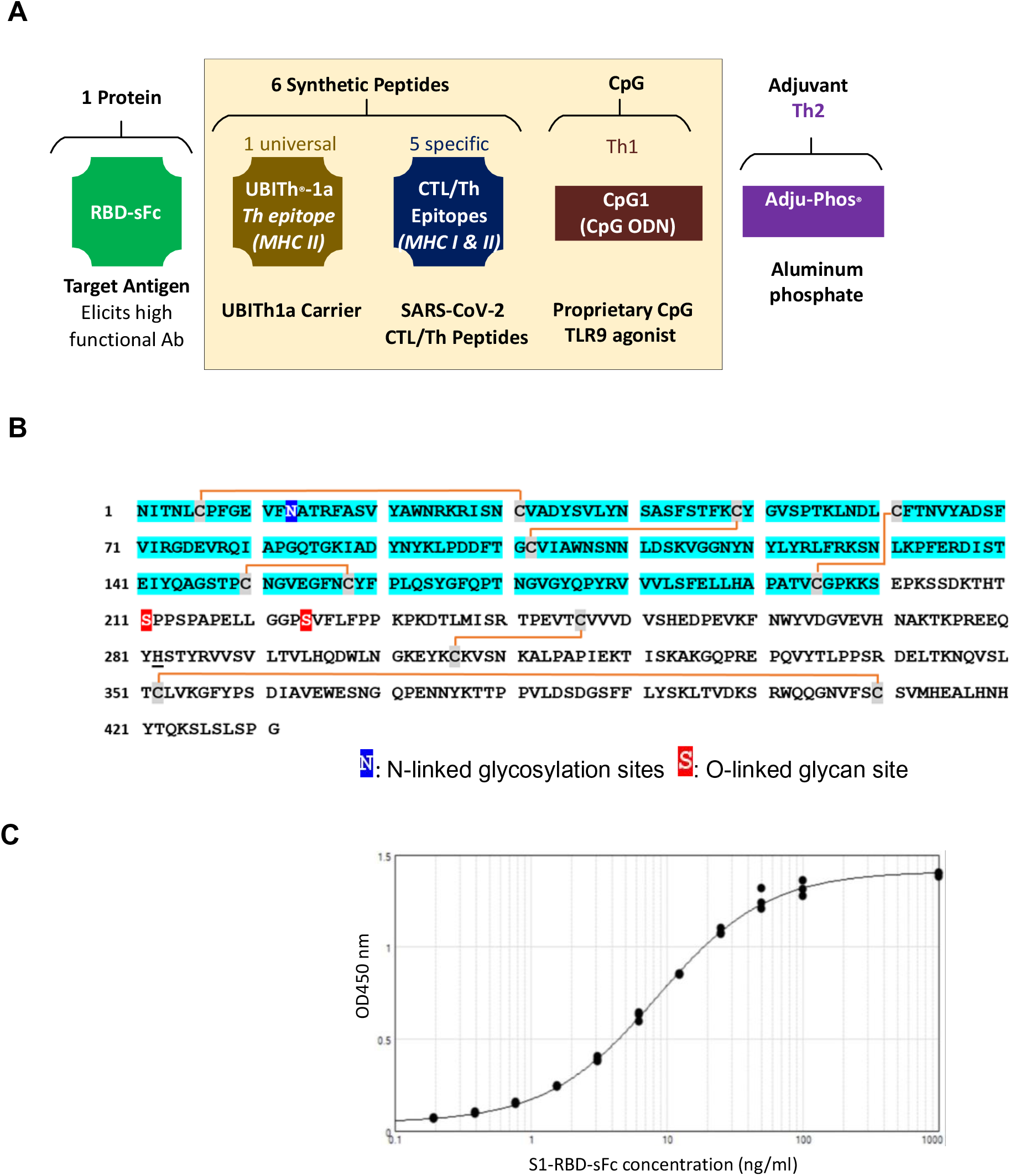
**Components and Sequence of the UB-612 Multitope Vaccine.** (A) UB-612 vaccine contains an S1-RBD-sFc fusion protein to elicit B cell responses, plus five synthetic Th/CTL peptides for class I and II MHC molecules derived from SARS-CoV2 S2, M, and N proteins, and the UBITh1a peptide (a proprietary T helper peptide). Among the 5 SARS-CoV2 peptides, there are 3 from S2, 1 from N and 1 from M proteins. These components are mixed with CpG1 and Adju-Phos adjuvant to constitute the UB-612 vaccine drug product. (B) Sequence of S1-RBD-sFc. S1-RBD-sFc protein is a glycoprotein consisting of one N-linked glycan (Asn13) and two O-linked glycans (Ser211 and Ser224). Light blue shading indicates RBD of SARS-CoV-2 and no shading indicates the sFc fragment of an IgG1. The substitution of His297 for Asn297 (EU-index numbering) in single chain Fc, His282 in S1- RBD-sFc, is indicated by underline. S1-RBD-sFc protein contains 431 amino acid residues, including 12 cysteine residues (Cys6, Cys31, Cys49, Cys61, Cys102, Cys150, Cys158, Cys195, Cys246, Cys306, Cys352 and Cys410), forming 6 pairs of disulfide bonds (Cys6- Cys31, Cys49-Cys102, Cys61-Cys195, Cys150-Cys158, Cys246-Cys306 and Cys352- Cys410), which are shown as orange lines. (C) hACE2 binding ability of S1-RBD-sFc, as determined via ELISA.

After *in vivo* identification of S1-RBD-sFc as the lead candidate, a stable cell line was generated through transfection of CHO cells followed by dihydrofolate reductase (DHFR) amplification. See methods section for full details of S1-RBD-sFc protein expression and purification.

### Modification of human IgG Fc portion

To reduce reactogenicity of S1-RBD-sFc, we modified the human IgG Fc portion of the protein. S1-RBD-sFc consists of the RBD linked with a human IgG1 sFc at the C-terminus (**Fig 1A**). The RBD domain functions as a high-affinity ligand for human Angiotensin-Converting Enzyme 2 (hACE2) cell receptors [37]. In this vaccine candidate, the IgG1 sFc domain was engineered to contain a series of mutations (C220S, C226S, C229S, and N297H), to eliminate the disulfide bonds and N-glycan, respectively (**Fig 1B**).

The mutation of N297 of the deimmunized heavy chain to H was to remove its glycosylation motif to prevent the depletion of target hACE2 through effector functions. S1-RBD-sFc was found to bind hACE2 well via ELISA (**Fig 1C**).

### Immunogenicity Study in Guinea Pigs to Down-Select S1- RBD-based-protein Design

The goal of the guinea pig (GP) immunogenicity study was to down select a single protein construct for use as the vaccine candidate. Three groups of GPs (N=5/group) were vaccinated at 0 and 3 weeks post initial immunization (WPI) intramuscularly (IM) with one of three S1-RBD-based protein immunogens formulated with ISA50 V2 adjuvant (**Fig 2A**). Sera were drawn at three time points (0, 3, and 5 WPI), and tested for immunogenicity through measurement of binding antibodies (BAbs) by ELISA. Inhibition of SARS-CoV-2 binding to hACE2 by RBD-Fc-elicited antibodies was investigated using ELISA and a cell-based assay.

**Fig 2.**
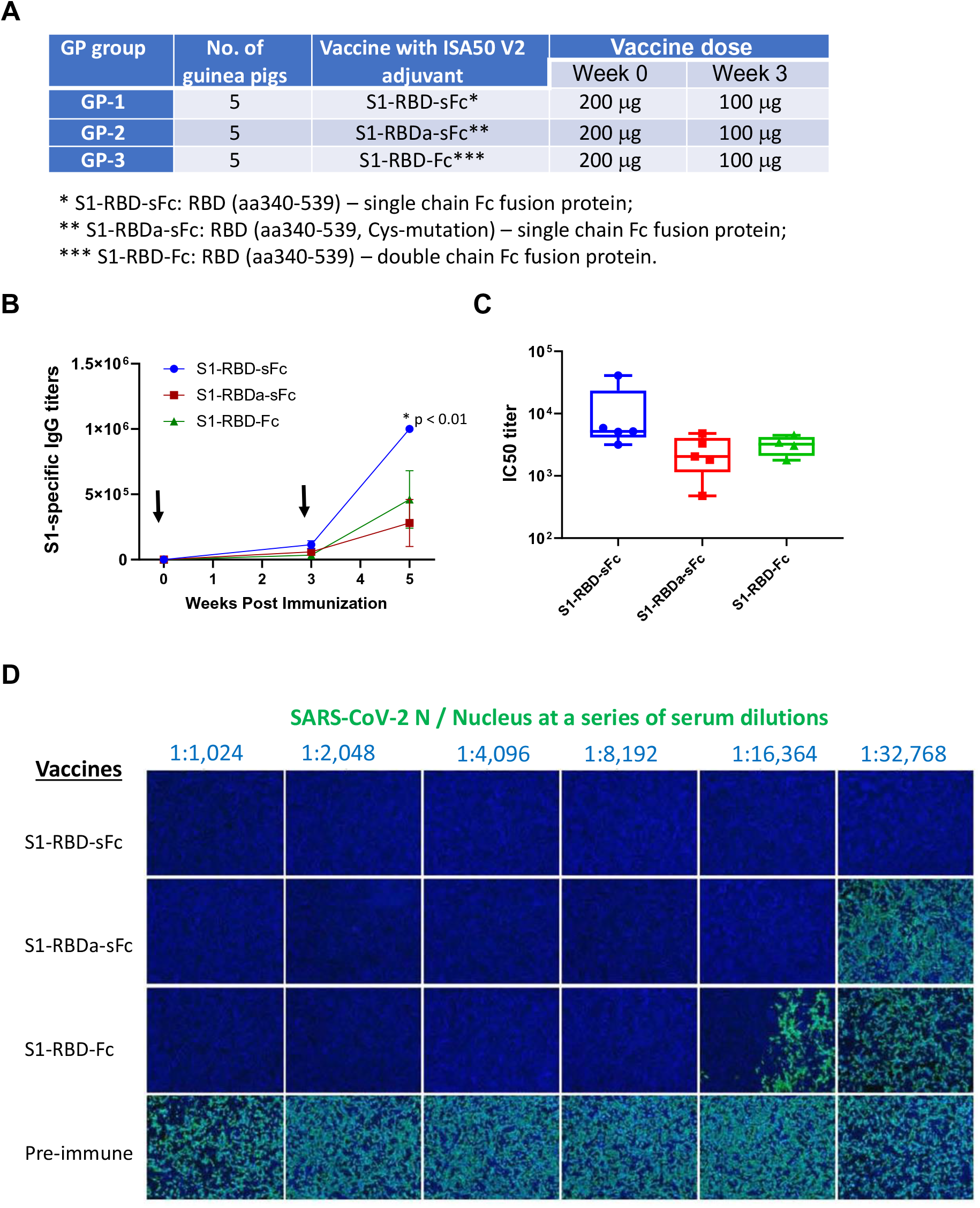
**S1-specific binding and neutralizing antibody responses of guinea pig immune sera.** (A) Guinea pig study design: animals were immunized with S1-RBD-sFC, S1-RBDa-sFC, or S1- RBD-Fc (n = 5 each group) at weeks 0 and 3 via intramuscular route. Immune sera were collected at 0, 3, and 5 weeks post initial immunization (WPI). Anti-S1 binding antibodies were detected by ELISA and neutralizing antibody titers were detected via CPE assay against wild type of SARS-CoV-2 virus. (B) S1-specific antibody temporal responses. Results shown as group geometric mean (GMT) ± standard error (SE). Immunization time points are shown with black arrows. * indicates that the statistical significance compared S1-RBD-sFC with S1-RBDa-sFC or S1-RBD-Fc immunization groups. (C) Neutralization and inhibitory dilution ID50 titers in S1 protein binding to ACE2 on ELISA by guinea pig sera collected at 2 weeks after the 2nd immunization (5 WPI). (D) Examples of nuclei count staining of Vero-E6 cells in the neutralization assay. Monolayers of Vero-E6 cells infected with virus-serum mixtures were assessed by immunofluorescence (IFA). Cells were stained with human anti-SARS-CoV-2 N protein antibody and detected with anti- human IgG-488 (green). The nuclei were counterstained with DAPI (4’,6-diamidino-2- phenylindole) (blue).

In the ELISA, recombinant SARS-CoV-2 spike protein S1 antigen was coated onto plates, and individual sera were tested for binding antibody titers to the coating antigen. As shown in **Fig 2A**, all constructs elicited a binding antibody response to SARS-CoV-2 S1 RBD protein in sera collected at 3- or 5-weeks post initial immunization (WPI). Of the three constructs tested, S1-RBD-sFc induced the highest immune response, with a geometric mean titer (GMT) nearly 5 log_10_ at 3 WPI and 6 log_10_ at 5 WPI. The difference between S1-RBD-sFc and S1-RBDa-sFc at 5 WPI was statistically significant (p ≦ 0.05), indicating that all constructs were highly immunogenic with S1-RBD-sFc holding an advantage in terms of binding antibody responses. For the S1:ACE2 binding inhibition activity evaluation, recombinant ACE2-ECD-sFc was immobilized onto plates. Individual serum was pre-incubated with His-tagged S1 protein (tracer) and then transferred into ACE2-ECD-sFc coated plates to test its inhibition activity. Although the mean ID_50_ inhibition values were not statistically significant (p ≥ 0.05), with the highest ID_50_ values observed for antibodies raised by S1-RBD-sFc (7251.5), followed by S1-RBD- Fc (3019.8) and S1-RBDa-sFc (1950.8). This result indicates that all antigens elicited antibodies capable of inhibiting hACE2 binding, with S1-RBD-sFc raising the most potent responses.

The function of anti-RBD antibodies was quantified both as inhibition of RBD binding to hACE2 and as neutralization of live SARS-CoV-2. In the hACE2 binding inhibition cell-based assay, HEK293 cells expressing hACE2 were treated with mixtures of pooled GP sera and S1-protein (Fc tagged), then assayed by flow cytometry (FACS) by staining cells with fluorescently labeled anti-human IgG Fc protein antibody. The GMT ID_50_ (the inhibitory dilutions at which 50% neutralization is attained) values were 1026 for S1-RBD-sFc, 193 for S1-RBDa-sFc, and 325 for S1-RBD-Fc vaccine, again showing functional activity for antibodies elicited by all candidates, still the highest activity was seen for S1-RBD-sFc. Results are provided in **Figs 2C**, **S1 and S2**, respectively. To test neutralizing capacity of the elicited antibodies, live virus cytopathic effect (CPE) 50% reduction assay were adopted, using the anti-SARS-CoV-2 N protein antibody and immunofluorescent visualization for neutralization titer (VNT_100_) determination (**Fig 2D and Fig S2**). Sera from S1-RBD-sFc demonstrated superior activity, with neutralization titers at 5 WPI 2-4-fold higher than those from the other two groups, protecting 50% of the cells from viral infection at titers of 504-1,024 at 3 WPI and >32768 in pooled guinea pig sera at 5 WPI (**Fig S2**).

In a separate experiment, we compared neutralizing titers in sera from GPs vaccinated with S1-RBD-sFc with convalescent sera of COVID-19 patients, using the S1- RBD:ACE2 binding inhibition ELISA (also termed as qNeu ELISA).The results, given in **Fig S2**, demonstrated that GP immune sera diluted 1,000-fold (3 WPI) or 8,000-fold (5 WPI) exhibited comparable or higher inhibition of S1-RBD:ACE2 binding than by the convalescent sera of 10 patients diluted at 20-fold, illustrating that the sera of GPs contained ≥50-fold higher antibody titers than human convalescent sera.

### Immunogenicity Studies in Rats

The initial immunogenicity assessment in GPs established the superior humoral immunogenicity of S1-RBD-sFc as the B cell component of our vaccine against SARS- CoV2. The GP experiments were tested with three protein candidates with a fixed dosing regimen (200 µg prime, 100 µg boost, ISA 50 adjuvant), allowing for a rigorous comparison of the respective candidate constructs. In the second set of experiments in Sprague-Dawley rats, the immunogen doses and adjuvants were varied to allow selection of an optimal adjuvant (**Fig 3A**). S1-RBD-sFc was formulated with five Th/CTL peptides selected from S2, M and N proteins of SARS-CoV-2 and our proprietary universal Th peptide (UBITh®1a) [32] to generate the multitope protein-peptide vaccine candidate (**Fig 1A**). We then combined the candidate vaccine with one of two different adjuvant systems: ISA51/CpG3 or Adju-Phos®/CpG1. These vaccine-adjuvant combinations were administered to rats IM on Week 0 (prime) and 2 (boost) with a wide dose range of 10 to 100 μg per injection. The animals were bled at baseline (day 0), 2 weeks., after 1st dose), 3 and 4 weeks (i.e., 1 and 2 weeks after the 2^nd^ dose) for antibody titer analyses.

**Fig 3.**
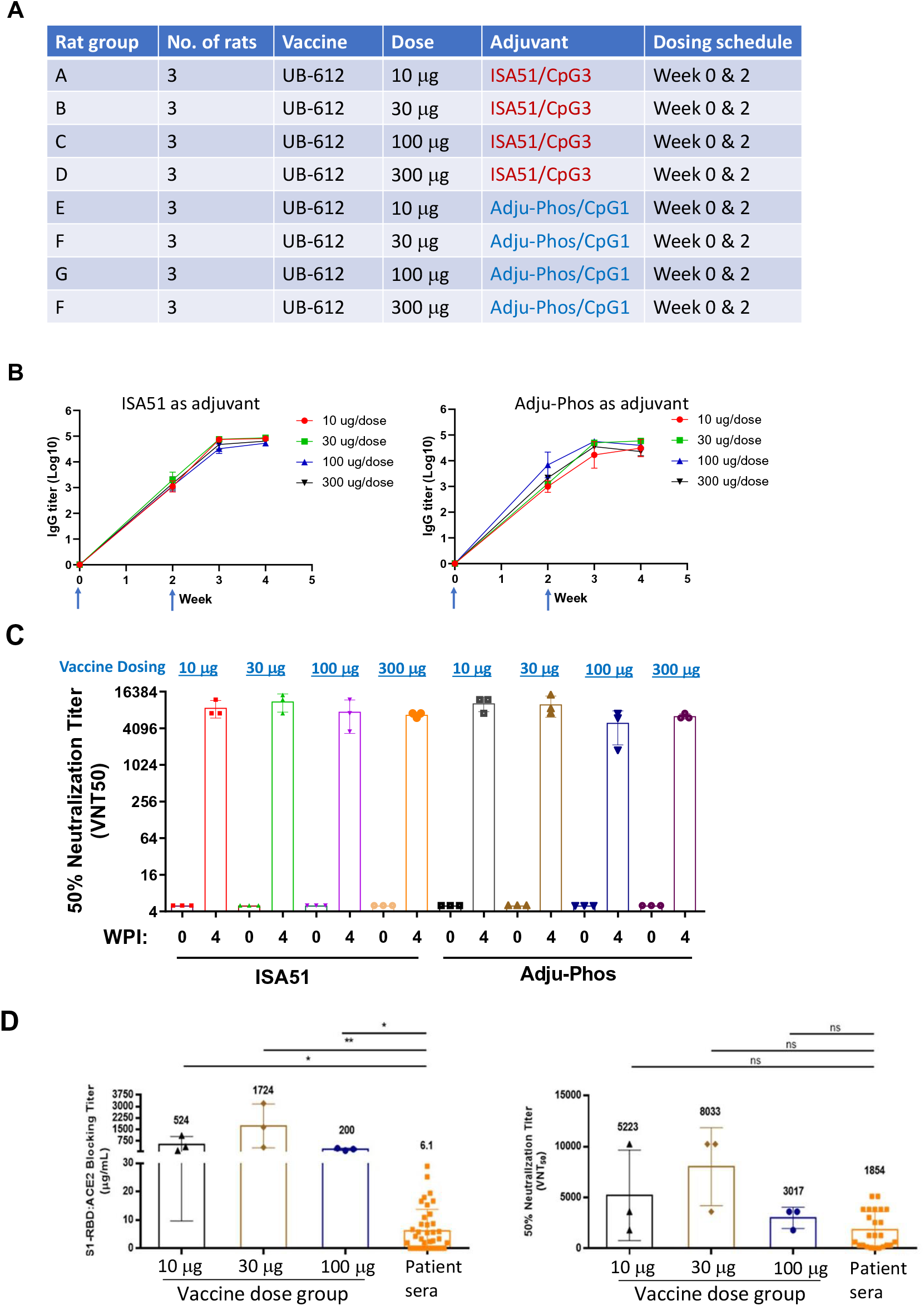
**Humoral immunogenicity testing in rats.** (A) Rat immunization study design. Immunogenicity of UB-612 adjuvanted with ISA51/CpG3 or Adju- Phos(R)/CpG1. Sprague Dawley rats were immunized at weeks 0 and 2 with UB-612 vaccine (at a dose range of 10-300 μg of S1-RBD-sF, formulated with synthetic designer peptides and adjuvants). (B) S1-RBD specific temporal antibody responses in immune sera at 0, 2, 3, and 4 WPI were assayed by ELISA. The rat groups in the left or right panels received vaccines with ISA51/CpG3 or Adju- Phos(R)/CpG1 as adjuvant. (C) Samples taken 4 WPI from rats immunized at weeks 0 and 2 with UB-612 vaccine adjuvanted with ISA51/CpG3 or Adju-Phos/CpG1. Potent neutralization of live SARS-CoV-2 by rat immune sera. Neutralization titers expressed as VNT50. (D) hACE2 binding inhibiting (left) and neutralizing antibody (right) titers of sera from UB-612 with Adju- Phos(R)/CpG1 vaccinated rats are higher than titers in convalescent COVID-19 patients (HCS). * *p* ≦ 0.05, ** *p* ≦ 0.01 and NS (not significant) (Kruskal- Wallis ANOVA with Dunn’s multiple comparisons test)

Vaccines formulated with either adjuvant system elicited similar levels of anti S1- RBD ELISA titers across all doses ranging from 10 to 100 µg, indicative of an excellent immunogenicity of the vaccine formulations even with low quantities of the primary protein immunogen (**Fig 3B**). In the S1-RBD:ACE2 binding inhibition ELISA, low doses of 10 and 30 µg induced inhibitory activity equivalent to the higher doses of 100 µg at Week 4. The most potent inhibitory activity was seen with the lowest dose of S1-RBD-sFc protein (10 µg) formulated with peptides and the Adju-Phos® adjuvant. In the replicating virus neutralization assay against the Taiwanese SARS-CoV-2 isolate (representative of the original Wuhan sequence), the Week 4 immune sera induced by UB-612 vaccine did not show a significant dose-dependent effect in rats (**Fig 3C**). The low doses of adjuvanted protein,10 and 30 μg, could neutralize viral infection at VNT_50_ of >10,240 dilution.

The rat immune sera at Week 6 (i.e. 4 weeks after the 2^nd^ immunization) from each vaccinated dose group were assayed two ways: first, in comparison with a set of convalescent sera of COVID-19 patients for titers in S1-RBD:ACE2 binding inhibition ELISA, expressed in blocking level of μg/mL; and second, through a SARS-CoV-2 CPE assay in Vero-E6 cells, expressed as VNT_50_. As shown in **Fig 3D**, all doses of the vaccine formulations elicited neutralizing titers in rats that were significantly higher than those in convalescent patients by S1-RBD:ACE2 binding ELISA and higher (but not achieving statistical significance due to variation in patient data and low number of animals) by VNT_50_.

To assess the Th1/Th2 response, vaccinated rats were evaluated using ELISpot. Rats were dosed at Weeks 0 and 2 with 30 µg or 100 µg of UB-612 vaccine. Splenocytes were then collected at Week 4 and restimulated *in vitro* with the Th/CTL peptide pool plus S1-RBD or with the Th/CTL peptide pool alone. High levels of IFN-γ and IL-2 secretion was observed in splenocytes after the stimulations with Th/CTL peptide pool plus S1-RBD or with the Th/CTL peptide pool alone, while only minor amounts of IL-4 were seen (**Figs 4A and 4B**). The individual peptide stimulations also demonstrated that the rat splenocytes also produced high levels of IFN-γ and IL-2 (Th1) responses but very low levels of IL-4 (Th2) against S2 peptides (p5752, p5753 and p5755) (**Fig 4S-A, B and C**); N peptide (p5754) (**Fig 4S-D**); panel E: M peptide (p5815) (**Fig 4S-E**). The results indicate that UB-612 is highly immunogenic and induces a Th1-prone cellular immune response, as shown by the high ratios of IFN-γ/IL-4 or IL-2/IL-4 [39].

**Fig 4.**
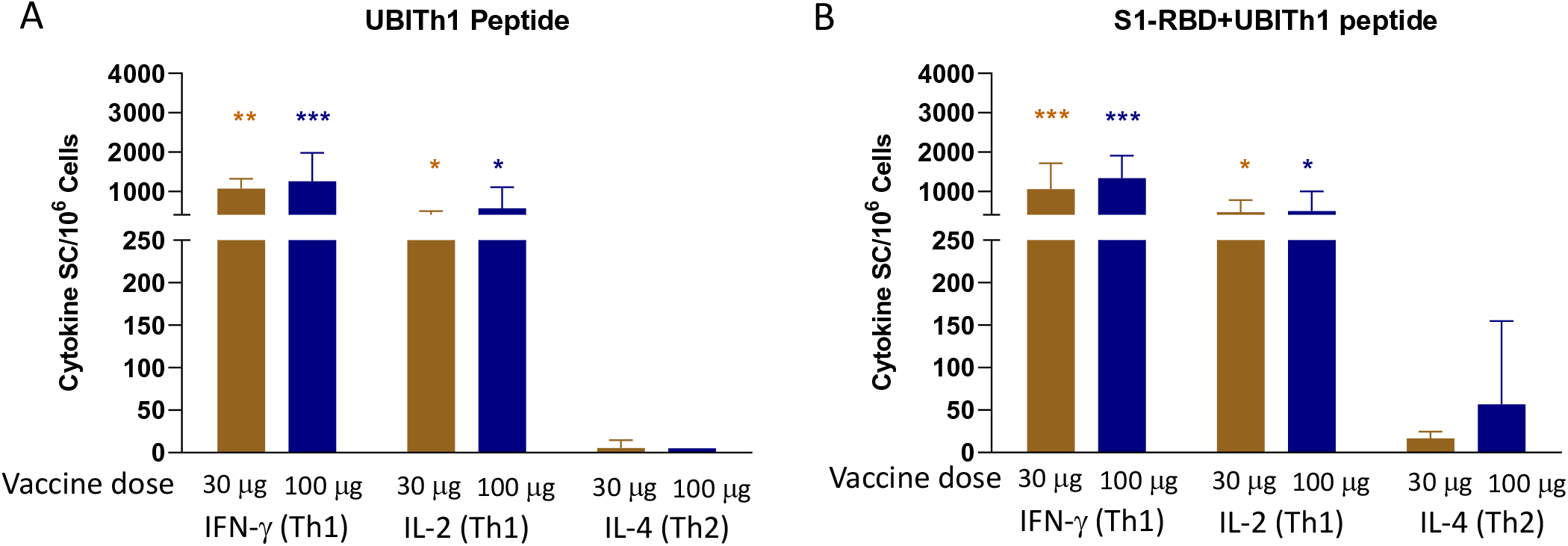
Cellular immunogenicity testing in rats (ELISpot detection of IFN-γ, IL-2 and IL-4 secreting cells in UB-612 immunized rats). Groups of rats were immunized with 30 μg or 100 μg UB-612 at Weeks 0 and 2. Splenocytes were collected at Week 4 and stimulated with the Th/CTL peptide pool alone (UBITh1 peptide, A) or with Th/CTL peptide pool plus S1-RBD (S1-RBD+UBITh1 peptide, B) used in the UB-612 vaccine composition. IFN-γ, IL-2 and IL-4-secreting splenocytes were determined by ELISpot analysis. Cytokine-secreting cells (SC) per million cells was calculated by subtracting the negative control wells. Bars represent the mean SD (n = 3). The secretion of IFN-γ or IL-2 was observed to be significantly higher than that of IL-4 in the 30 and 100 µg group (* *p* < 0.05, ** *p* < 0.01, *** *p* < 0.005). (A) ELISpot detection of IFN-γ, IL-2 and IL-4 responses from cells stimulated with UBITh1 peptide pool. (B) ELISpot detection of IFN-γ, IL-2 and IL-4 responses from cells stimulated with UBITh1 peptide pool in combination with S1-RBD

### Protective immunity in AAV-hACE2 transduced mice

The initial challenge study of UB-612 was performed in the adeno-associated virus (AAV)/hACE2 transduced BALB/c mouse model. Groups of 3 BALB/C mice were vaccinated at Weeks 0 and 2 with UB-612 containing 3, 9 or 30 µg of protein and formulated with Adju-Phos®. S1-RBD-specific antibody titers were evaluated at Weeks 0, 3 and 4. After 2 doses of vaccine, S1-RBD specific antibody responses were detected in all three dose groups with significant dose dependent response pattern (**Fig 5B**), *p* < 0.05 between 3 and 9 µg groups and *p* < 0.005 between 3 and 30 µg groups. The mice were infected with adeno-associated virus (AAV) expressing hACE2 at 4 WPI and challenged 2 weeks later with 10^6^ TCID_50_ of SARS-CoV-2 by the intranasal (IN) route (**Fig 5A**). Efficacy of the vaccine was measured using lung viral loads and body weight measurements. As shown in **Fig 5C**, vaccination with 30 μg of UB-612 significantly reduced lung viral loads (∼3.5 log_10_ viral genome copies/ug RNA or ∼ 5-fold TCID_50_/mL of infectious virus) compared to the saline group (p <0.05, paired t-test). As shown in **Fig 5D**, vaccination with middle (9 μg) and high (30 μg) doses resulted in a reduction in lung pathology. There was no evidence even in the suboptimal (3 μg) dose group of enhancement of lung pathology (**Fig 5F**). The lung pathological scores shown in **Fig 5E** demonstrated that the high dose (30 μg) group had significant pathological score reduction compared to the saline group, *p* < 0.05. Vaccination with 3 or 9 µg of UB-612 reduced live virus detection by cell culture method (TCID_50_) to below the level of detection (LOD) but did not appear to reduce viral loads significantly when measured by RT-PCR. In sum, despite the lack of statistical power (N=3 mice) in this study, it appears that the highest dose of 30 µg could have maximum protective efficacy as demonstrated by the absence of live virus, inflammatory cell infiltration, and immunopathology in the lungs.

**Fig 5.**
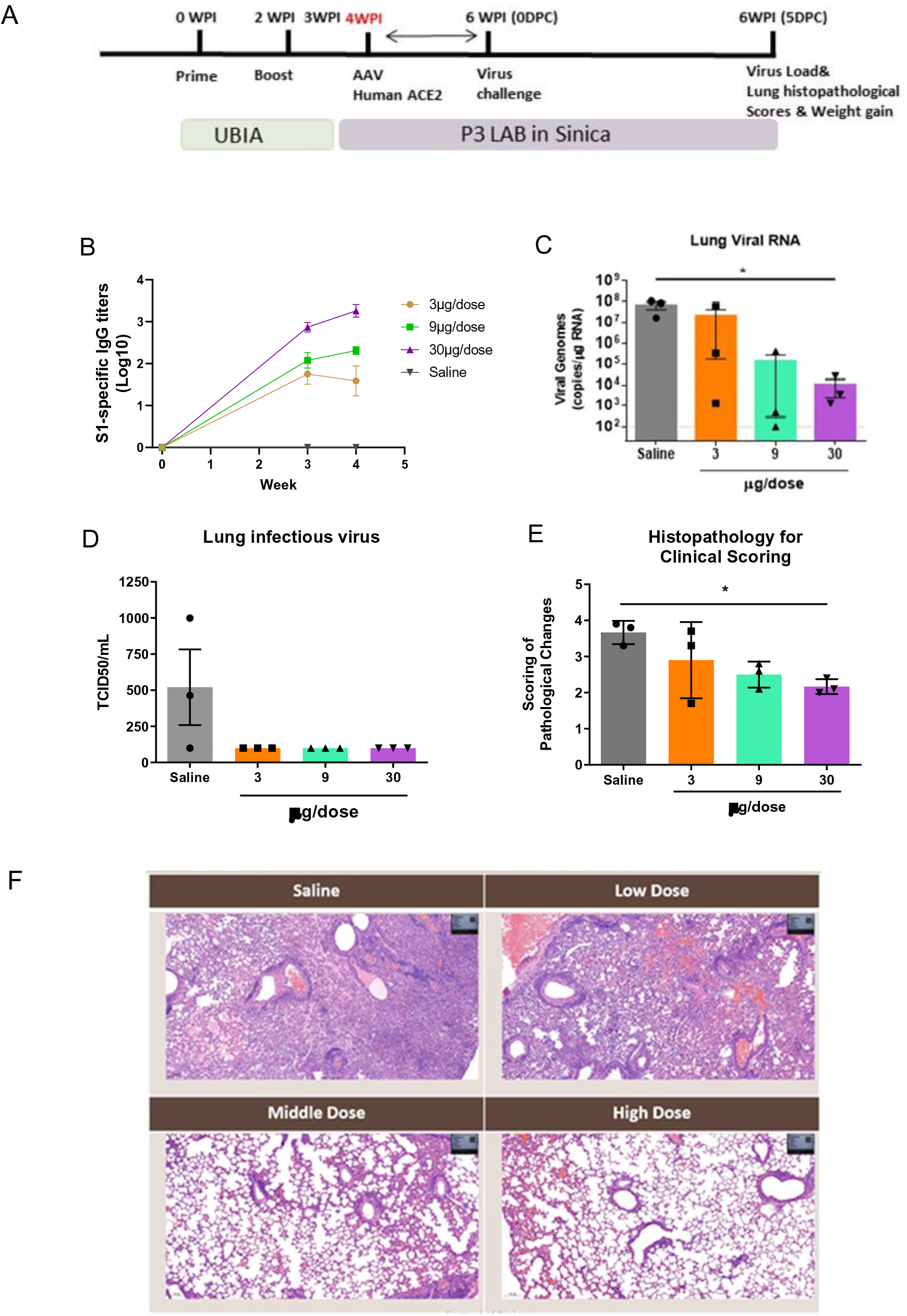
UB-612 vaccine immunogenicity in BALB/C mice and protective immunity against live SARS- CoV-2 challenge in hACE-transduced mice. Groups of BALB/C mice were immunized with 3 μg, 9 μg or 30 μg UB-612 vaccine or saline at Weeks 0 and 2. Serum samples collected at Weeks 3 and 4 were evaluated for S1-RBD-specific antibody responses. The mice were transduced with AAV-RBD at Week 4 and challenged with 10^6^ PFU TCID_50_ of SARS-CoV-2 virus (hCoV-19/Taiwan/4/2020) by intratracheal infection at Week 6. The lung viral load and pathology were detected at 5 days after infection. (A) Immunization and challenge schedule. (B) S1-RBD-specific antibody titers at Weeks 0, 3 and 4 were measured with significant dose dependent response trend. * *p* < 0.05 between 3 and 9 µg groups; *** *p* < 0.005 between 3 and 30 µg groups. (C) SARS-CoV-2 viral load RNA in lung were determined by RT-PCR. Significant difference is indicated between the saline and 30 µg groups, * *p* < 0.05. (D) The live virus titers in lung were determined by TCID_50_. (E) Lung pathological scores on Day 5 after challenge. Significant difference is indicated between the saline and 30 µg groups, * *p* < 0.05. (F) Lung pathology. Stained sections of mouse lung tissues from different vaccination groups of mice challenged with live virus. The vaccine dose: Low dose: 3 μg; Middle dose: 9 μg, or High dose: 30 μg UB-612 vaccine; and Saline as negative control.

### Immunogenicity and challenge studies in rhesus and cynomolgus macaques

Based on an established model using rhesus macaques (RM) [37, 38], an immunization study of UB-612 by IM injection was initiated with RM (N = 4/group) receiving 0, 10, 30 or 100 μg of UB-612 at 0 and 4 weeks in the first NHP study (Study 1) (**Fig 6A**). IgG binding antibody to S1-RBD was increased over baseline in all animals, with titers reaching around 3 logs at 5 and 7 weeks (**Fig 6B**). Strong neutralizing antibody responses were induced, with highest titers observed at the 30 μg dose (**Fig 6C**). In ELISpot antigen-specific IFN-γ-secreting T cells were elicited in a dose-dependent manner (**Fig 6D**), with highest responses at the 100 μg dose level. To test the response to boosting, the 3^rd^ immunization was given at Day 70 (6 weeks after the 2^nd^ immunization). One week after the 3^rd^ dose, S1-specific IgG titers were significantly boosted (∼5-fold) at all three dose levels (**Fig 7A**). Neutralizing antibody responses against the Wuhan strain also increased in a live virus CPE assay one week after the 3^rd^ dose, with the greatest increase (5∼10- fold) seen for the 100 μg dose level (**Fig 7B**). We also measured neutralization in a pseudovirus assay expressing the Spike proteins from the Wuhan strain and 5 variants of concern (VOC: B.1.1.7, P.1, B.1.429, B.1.526 and B.1.351) on Days 42 and 70 after initial immunization and one week after the 3^rd^ dose. Serum collected one week post third dose had increased neutralization activity against all five VOC (**Fig 7C**). In a live virus assay, sera taken one week after the 3^rd^ dose (Day 77) demonstrated potent neutralization responses against the D614G strain and four variants of concern. (**Fig 7D**). The neutralizing activities against Wuhan (wild-type, WT), D614G, as well as Variant of

**Fig 6.**
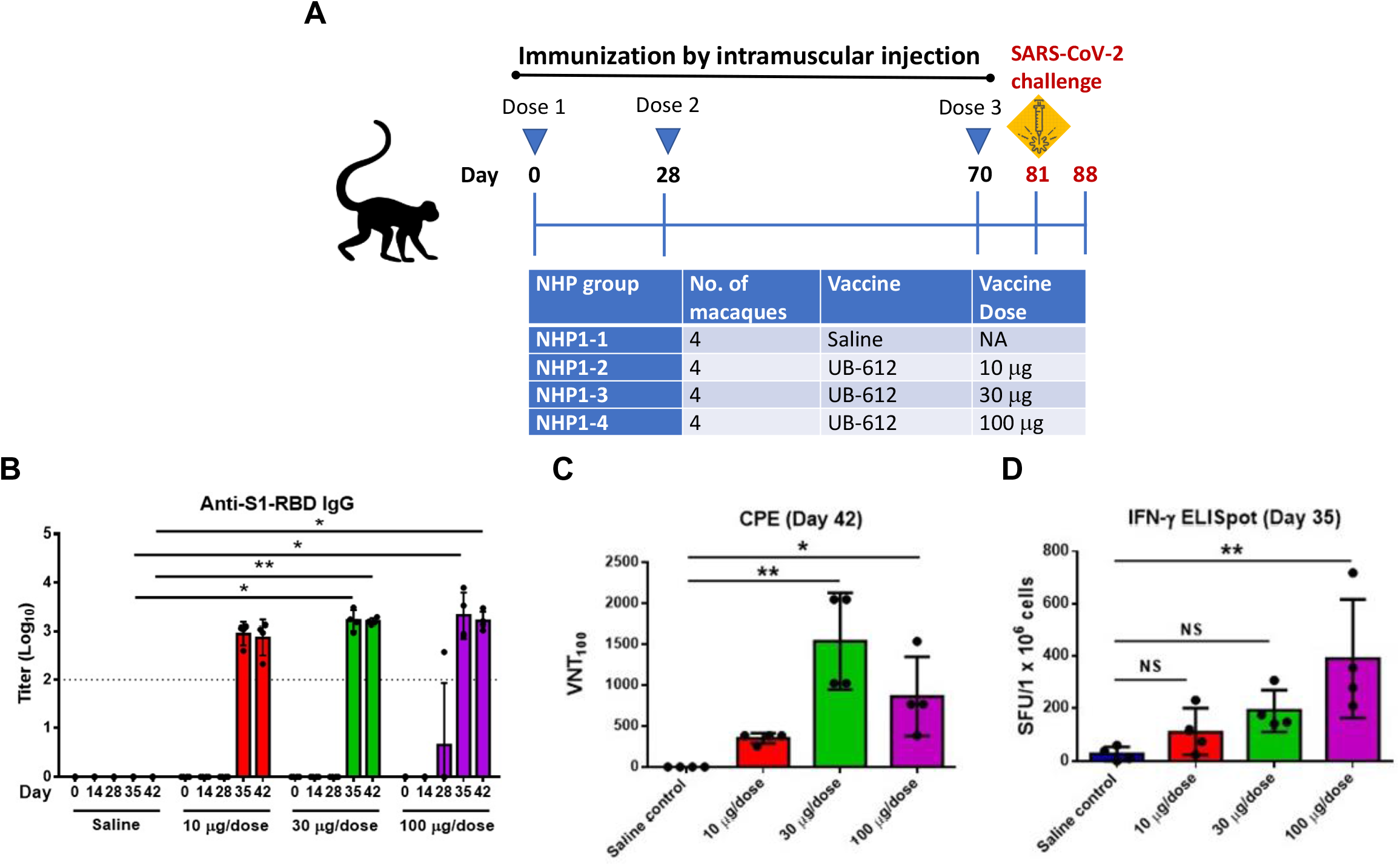
**Immunogenicity results in rhesus macaques (Study 1).** (A) The immunization study design and groups in rhesus macaques (Study 1). (B) Direct binding of rhesus macaque (RM) immune sera to S1-RBD on ELISA. ELISA-based serum antibody titer (mean Log_10_ SD) was defined as the highest dilution fold with OD_450_ value above the cutoff value. * *p* ≦ 0.05, ** *p* ≦ 0.01 (C) Potent neutralization of live SARS-CoV-2 by RM immune sera. Immune sera collected at Day 42 from RM vaccinated at weeks 0 and 4 were assayed in SARS-CoV-2 infected Vero-E6 cells for cytopathic effect (CPE). (D) ELISpot analysis of RM PBMC cells stimulated with Th/CTL peptide pool. PBMCs were collected at Day 35 and stimulated with Th/CTL peptide pool. IFN-γ-secreting cells were determined by ELISpot analysis. ** *p* ≦ 0.01.

**Fig 7.**
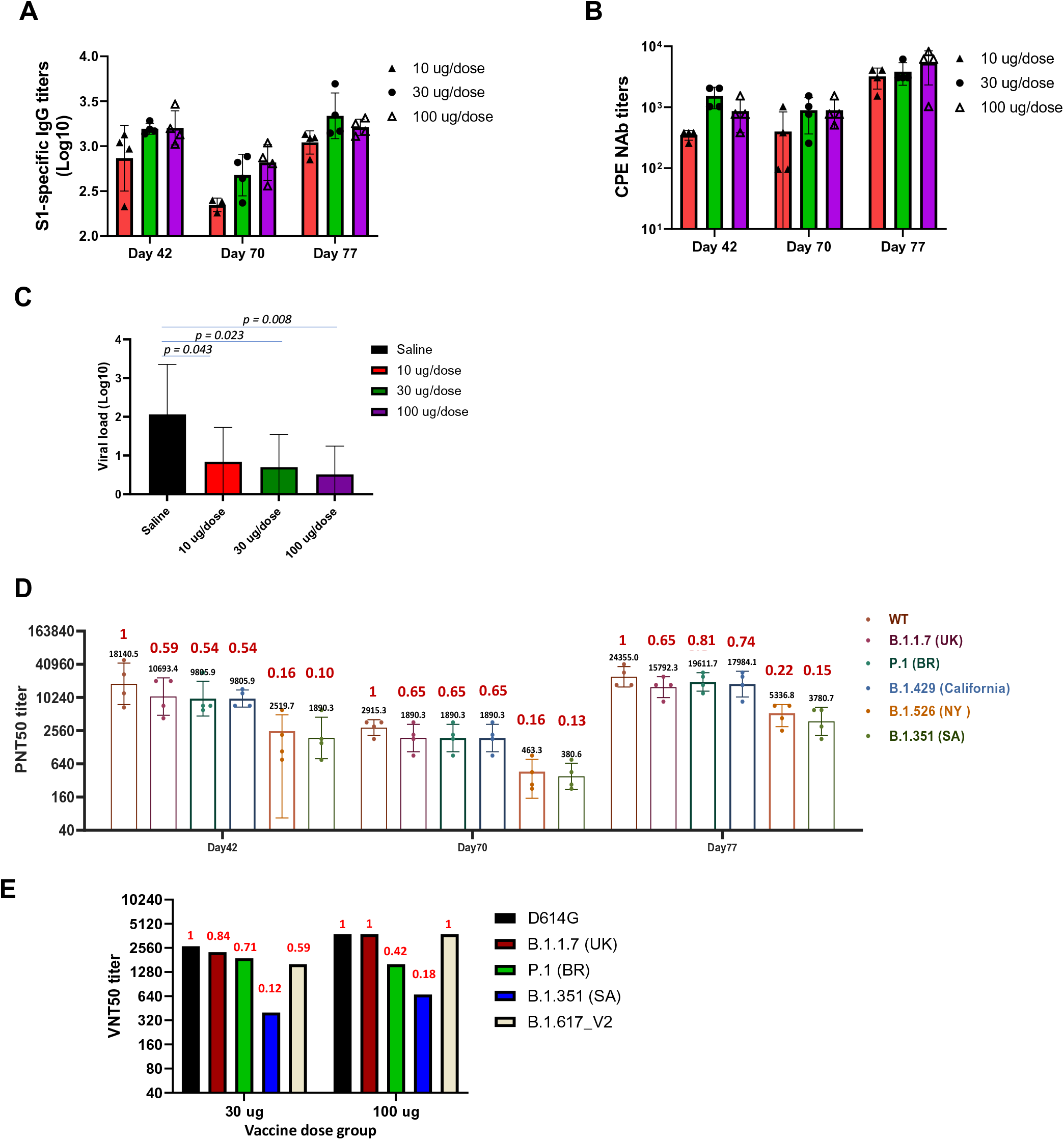
**The 3^rd^ dose of vaccine boosted S1-specific IgG titers and neutralizing antibody responses in rhesus macaque (Study 1) and significantly reduced viral load in lung after SARS-CoV-2 challenge.** (A) Direct binding of rhesus macaque (RM) immune sera to S1-RBD on ELISA in macaque serum samples collected on Day 42 (14 days after the 2nd immunization), Day 70 (prior to the 3^rd^ immunization), and Day 77 (1 week after the 3rd immunization). (B) Potent neutralization of live SARS-CoV-2 by RM immune sera by CPE assay in macaque serum samples collected on Day 42, Day 70, and Day 77. (C) The macaques were challenged at 11 days after the 3^rd^ immunization (Day 81) with SARS-CoV- 2 (10^6^ TCID50) intratracheally. The viral loads were determined by viral RNA copies/gram of lung tissue at 7 days after challenge (Day 88). Neutralizing antibody responses against SARS-CoV-2 pseudoviruses expressing the Spike proteins from wild type Wuhan strain (WT) and 5 variants of concerns (VOCs: B.1.1.7, P.1, B.1.429, B.1.526 and B.1.351) in macaque serum samples collected on Day 42, Day 70, and Day 77 from the 100 μg vaccine dose group. After the 3^rd^ immunization, the NAb titers were boosted. The numbers on each bar indicate the ratio of GMT VNT_50_ of VOC/GMT VNT_50_ of WT. (D) Neutralizing antibody responses against SARS-CoV-2 live viruses D614G and variants (B.1.1.7, P.1, B.1.429, B.1.351 and B.1.617_V2) in macaque serum samples collected from both 30 mg and 100 mg vaccine dose groups on Day 77 (7 days after the 3rd immunization). The numbers on each bar indicate the ratio of GMT VNT50 of VOC/GMT VNT50 of D614G.

Concerns (VOCs) B.1.1.7, P.1, B.1.429 and B.1.617 V2 were very similar while the neutralizing activities were reduced against B.1.526, B.1.351 by >50% compared to WT or D614G strains (**Fig 7D and 7E**). Animals were challenged on Day 77 with SARS-CoV-2 (strain and dose). Reduced nasal shedding and lung segment viral loads were significantly reduced (**Fig 7C**). There was no evidence for enhancement of lung pathology

In a second NHP study, cynomolgus macaques received either saline, UB-612 30 μg or UB-612 100 μg on Day 0 and Day 28 and were challenged intratracheally (IT) with 1x 10^5^ TCID50 SARS-CoV-2 virus (Wuhan strain) on Day 55 (**Fig 8A**). The results showed that high titers of neutralizing antibody titers were achieved against live virus Wuhan strain on Day 50 (3 weeks after the 2^nd^ immunization for both the 30 μg and 100 μg doses of UB-612) (**Fig 8B**) and also against the B.1.617.2 Delta variant (**Fig 8C**), though a drop in titer of 2-fold for the 30 μg dose and 1.5-fold for the 100 μg dose was seen for Delta as compared to the Wuhan strain. When viral loads were measured in the bronchoalveolar lavage (BAL), nasal swabs, and rectal swabs through subgenomic RNA RT-PCR, the 100 μg dose group was associated with the highest protective immunity (**Figs 8D-8F**).

**Fig 8.**
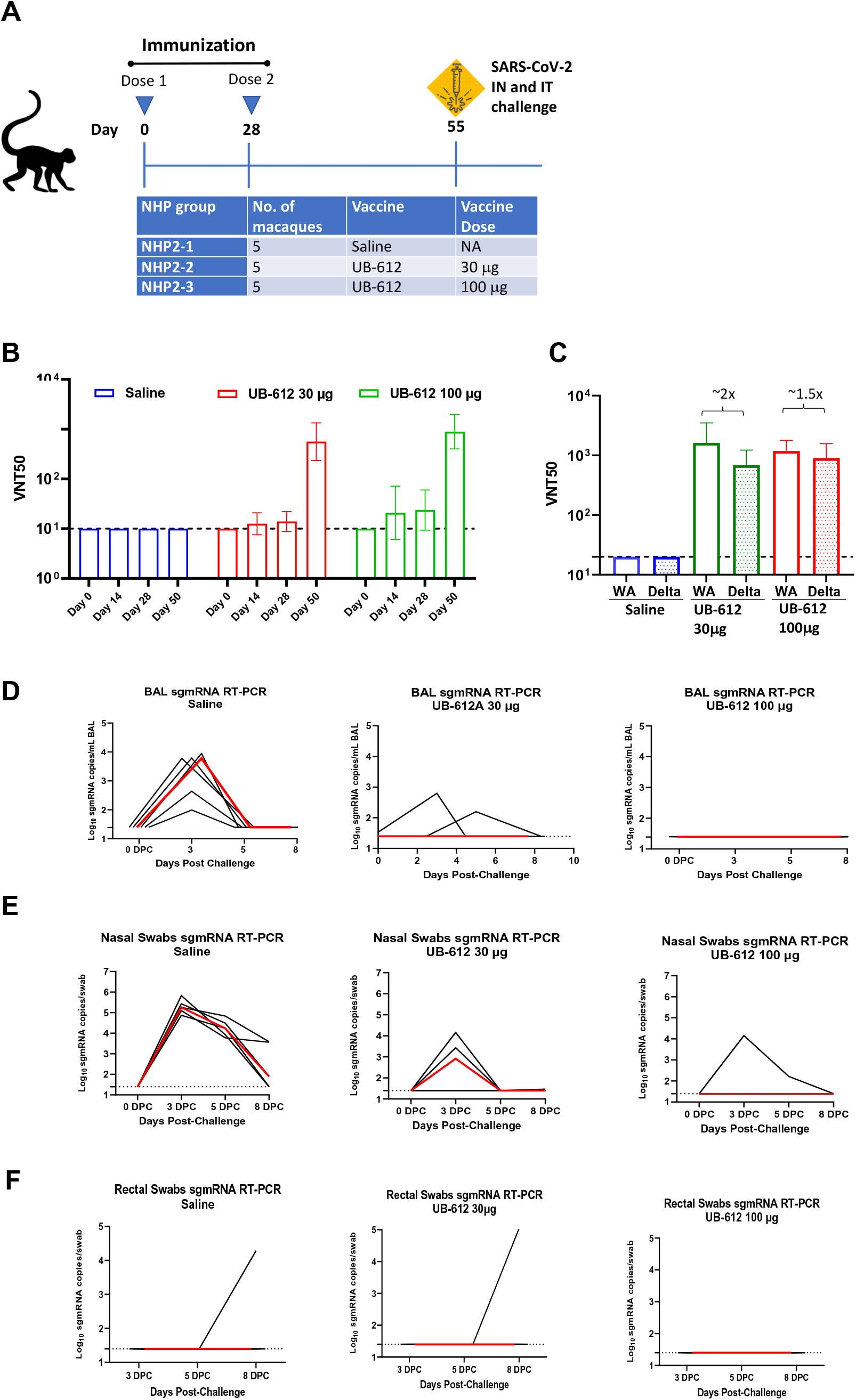
Neutralizing antibody response and protective immunity in cynomolgus macaques (Study 2). The macaques received Saline, UB-612 30 μg or UB-612 100 μg on Day 0 and Day 28, and were challenged with SARS-CoV-2 virus on Day 55. (A) The immunization and challenge study design in rhesus macaques (Study 2). SARS-CoV-2 Wuhan strain was used for challenge on Day 55 with a total of 1.0 × 105 TCID50 of SARS-CoV-2 divided equally between intranasal (IN, 0.5x 105 TCID50) and intratracheal (IT, 0.5x105 TCID50) administration. (B) Neutralizing antibody responses against wild type Wuhan (WA) strain at different time points. (C) Neutralizing antibody tiers against WA and Delta variant on Day 50 (3 weeks after the second immunization). (D-F) Viral loads were detected in (D) BAL, (E) nasal swabs and (F) rectal swabs through sgmRNA RT-PCR after challenge with SARS-CoV-2 Wuhan strain. The black curves represent viral loads of the individual animals, and the red curves are the median viral load of each group.

## Discussion

Successful vaccines against many viral diseases including COVID-19, protect through eliciting neutralizing antibody responses. However, although neutralizing antibodies may fully protect at titers present weeks after the initial primary vaccination, protection may wane over time. Under this circumstance, clonally expanded populations of antigen-specific lymphocytes may be important for maintaining protection. Falling neutralizing titers also raise the possibility of antibody-dependent enhancement (ADE) or concerns over Vaccine Associated Enhanced Respiratory Disease (VAERD) by SARS-CoV-2 [9,39,40]. UB-612 was developed specifically to address the need for a vaccine that elicits a strong, but ADE-free neutralizing response while also eliciting robust and long-lasting T cell responses. The ADE-free neutralizing response was accomplished by avoiding amino acid residues 597-603, located in the S2 subunit, which have been implicated in ADE of SARS-CoV *in vitro* and in NHPs [8].

Initially, we developed three RBD-sFc fusion protein vaccine candidates, which were then down-selected to a single candidate through immunogenicity tests in GPs, showing robust S1-RBD binding antibody responses and functional activity, including neutralization of live SARS-CoV-2 and inhibition of s1:hACE2 binding. Of the three candidates tested, S1-RBD-sFc (S1-RBD fused to a single-chain Fc) gave the strongest responses in all measurements. S1-RBD-sFc was slightly more immunogenic than the other two constructs in S1-RBD binding antibody assays, but the strength of the S1-RBD- sFc immunogen became abundantly clear when functional activity of elicited antibodies was tested. Functional activity was quantified both as inhibition of viral binding to hACE2 and as neutralization of live SARS-CoV-2. In both tests, all antigens elicited functional antibodies, with S1-RBD-sFc raising the most potent responses. Based on these results, the S1-RBD-sFc protein was selected as the lead candidate for the B cell component of the vaccine.

Further confirmation of the immunogenicity of this vaccine candidate was obtained by comparison of neutralizing titers in sera from S1-RBD-sFc-immunized GPs with titers in convalescent sera from COVID-19 patients. The results demonstrated that highly diluted GP immune sera (e.g. 1:1000 after one dose or 1:8000 after two doses) exhibited comparable or higher inhibition of S1-RBD:ACE2 binding than convalescent sera of 10 patients diluted 20-fold. This finding is of clinical significance since it suggests that an S1- RBD-sFc-based vaccine can elicit sufficient neutralizing antibodies to prevent SARS-CoV- 2 infection.

S1-RBD-sFc was then combined with rationally designed Th/CTL peptides, derived from S2, M and N structural proteins of SARS-CoV-2, to generate the final unadjuvanted vaccine candidate, which was studied in Sprague Dawley rats to compare two adjuvant combinations (ISA 51VG/CpG3 and Adju-Phos®/CpG1).

In the initial rat study, which tested the two vaccine-adjuvant combinations at a dose range of 10 to 300 μg per injection, our results indicated that vaccines formulated with both adjuvant systems elicited similar BAb titers across all doses, indicating excellent immunogenicity of the vaccine even for low quantities of the primary protein immunogen. As in the earlier GP studies, however, functional antibody assays demonstrated clear differences between candidate vaccine formulations. These tests showed the equivalency in immunogenicity between the two adjuvant combinations while confirming excellent immunogenicity at low doses. We chose Adju-Phos®/CpG1 as the adjuvant in our final vaccine formulation due to its long safety record and ability to improve immune responses. Neutralization results indicated that even low doses of adjuvanted protein (10 and 30 μg) showed excellent neutralizing immunogenicity. All doses of protein elicited neutralizing titers significantly higher than those in convalescent patients, as determined through an S1-RBD:ACE2 binding ELISA, Additionally, titers were higher (but not achieving statistical significance due to the spread in patient data and the low number of animals) by VNT_50_.

A Th1-oriented immune response against SARS-CoV-2 is potentially important to avoid ADE or VAERD, as demonstrated by studies with SARS and MERS coronaviruses as well as a commercial formalin inactivated RSV vaccine (inducing a Th2-biased response), which led to the death of several vaccinated children who were later exposed to live RSV [9,41,42]. Therefore, the FDA has recommended that any vaccines for COVID- 19 provide data of a Th1-biased immune response in animals before proceeding to First in Human (FIH) trials [43]. To address this issue, T cell immunity was assessed in rats upon restimulation of splenocytes harvested from vaccinated animals with the respective B and T cell vaccine components. Our results indicate that UB-612 vaccination can induce a robust Th1-prone cellular immune response, likely due to presence of CpG1 [44], even under the influence of a Th2-biased alum-containing adjuvant system [45].

We used an AAV/hACE2 transduced BALB/c mouse model developed by Dr. Tao, Mi-Hua at Academia Sinica in Taiwan [46] to demonstrate protective efficacy of UB-612 *in vivo*. Wild-type mice are not a suitable host for SARS-CoV-2; however, mice expressing hACE2 are susceptible to infection and disease from SARS-CoV-2 [47]. In this mouse model, productive infection with SARS-CoV-2 leads to high viral loads in the lungs and weight loss, which can be used together with other clinical symptoms to assess safety, (e.g., lack of ADE or VAERD) and efficacy of vaccine candidates including UB-612. We infected mice with AAV-expressing hACE2 at 4 WPI and challenged 2 weeks later with SARS-CoV-2 via the intranasal (IN) route. Our results demonstrate that UB-612 can reduce lung viral load and weight loss without induction of VAERD or immunopathology in the lungs in a dose-dependent manner.

In addition, we tested UB-612 in rhesus and cynomolgus macaque models. While SARS-CoV-2 does not cause a lethal COVID-19-like disease in monkeys, the virus can cause infection and illness. In these animals, disease is generally mild, self-limiting and resolves within 2 weeks [48–50]. In an initial study, rhesus macaques received three vaccinations at three dose levels. All vaccinated animals developed high titers of S1-RBD binding antibodies that were also potently neutralizing. In a second study, cynomolgus macaques were immunized with 2 IM administration of either 30 or 100 ug of UB-612 and challenged via the intratracheal and intranasal routes. All immunized animals demonstrated strong immune responses and were protected against challenge.

In summary, we have developed and demonstrated proof of concept for UB-612, a novel multitope protein-peptide vaccine being rapidly advanced in clinical trials for prevention of SARS-CoV-2 infection and COVID-19. We showed that our vaccine elicited high levels of neutralizing antibodies and a Th1-prone immune response that protected animals challenged with a high dose of SARS-CoV-2, without induction of immunopathology in the lungs.

## Materials and Methods

### Vaccine Design and Production

#### Design of UB-612 Vaccine Immunogen

The UB-612 vaccine immunogen was designed to contain an S1-RBD-sFc fusion protein plus five synthetic Th/CTL peptides for class I and II MHC molecules derived from SARS-CoV-2 S2, M, and N proteins. For Th/CTL epitope design, we employed the “Epitope Prediction and Analysis Tools” (http://tools.iedb.org/population) to identify the desirable CTL T cell epitopes. Th/CTL epitopes from highly conserved sequences derived from all three SARS-CoV-2 proteins (S, N and M) proteins were identified through an extensive literature search and epitope analysis [22, 29]. Five peptides within these regions were selected for inclusion in the UB-612 immunogen and subject to further designs. Each selected peptide contained Th or CTL epitopes with prior validation of MHC I or II binding [51] and exhibited good manufacturability characteristics (optimal length and amenability for high quality synthesis). These rationally designed Th/CTL peptides were further modified by addition of a Lys-Lys-Lys tail to each respective peptide’s N-terminus to improve peptide solubility and enrich positive charge for use in vaccine formulation [32]. UBITh®1a, a proprietary synthetic peptide with an original framework sequence derived from the measles virus fusion protein (MVF) was added to S1-RBD-sFc to enhance the immune response. Aluminum phosphate (Adju-Phos®, InvivoGen) adjuvant was also added to further promote Th2 responses. These components were mixed with CpG1, which binds the positively (designed) charged peptides by dipolar interactions and also serves as an adjuvant. All peptides and CpG1 were produced synthetically.

#### Vaccine Construction and Preparation

S1-RBD-sFc protein contains 431 amino acid residues with 12 cysteine residues (Cys6, Cys31, Cys49, Cys61, Cys102, Cys150, Cys158, Cys195, Cys246, Cys306, Cys352 and Cys410), forming 6 pairs of disulfide bonds (Cys6-Cys31, Cys49-Cys102, Cys61-Cys195, Cys150-Cys158, Cys246-Cys306 and Cys352-Cys410). The molecular mass of S1-RBD-sFc protein is about 50 kDa. To construct the vector expressing the recombinant fusion protein, the cDNA sequence encoding the S1-RBD-sFc protein (**Fig 1B**) was synthesized, digested, and then ligated into Freedom® pCHO 1.0 vector (Life Technology) to obtain the pCHO S1-RBD-sFc expression vector. The cDNA sequence was confirmed by DNA sequencing. The S1-RBD-sFc protein was expressed in transient transfections of CHO cells for GP and rat immunogenicity studies. Three forms of cDNA fragments for S1-RBD-sFc, S1-RBDa-sFc and S1-RBD-Fc fusion proteins were designed for transient expression in ExpiCHO-S system for target protein production. The cell culture was harvested 12–14 days post-transfection, clarified by centrifugation and 0.22- μm filtration, and purified by protein A chromatography. The purity of the fusion proteins was determined on SDS gel, and protein concentration was determined according to the optical density (OD) of UV absorbance at a wavelength of 280 nm. To formulate S1-RBD- based protein vaccines at 200 µg/mL, equal volumes of the immunogen (S1-RBD sFc, S1-RBDa-sFc or S1-RBD-Fc) in aqueous phase at 400 µg/mL and sterile water-in-oil adjuvant MontanideTM ISA50 V2 (Seppic) were mixed using two syringes attached to a 3-way Stopcock and emulsified until no phase separation was seen. The emulsified vaccines were stored at 4°C before use.

#### Generation of stable cell line for S1-RBD-sFc and confirmation of binding activity

A stable high-expressing clone was isolated, and a cell bank was produced using standard stable CHO methods. S1-RBD-sFc was produced in a suspension culture, purified by multi-step column chromatography and characterized. Peptide mapping, N- and C-terminal amino acid sequencing, and analysis of disulfide bonding and glycosylation confirmed that the expressed and purified protein conformed to the predicted characteristics. Size-exclusion chromatography (SEC), analytical ultracentrifugation, and capillary electrophoresis with sodium dodecyl sulfate (CE-SDS) experiments demonstrated that S1-RBD-Fc exists in two major isoforms, S1-RBD-sFc1 and S1-RBD- sFc2, corresponding to N-linked and O-linked glycoforms of the protein. The binding activity of the vaccine was tested in an hACE2 ELISA and was demonstrated to bind hACE2 with an EC50 of 8.477 ng/mL, indicative of high affinity.

### Animal Procedures for Immunogenicity Studies

#### Immunogenicity studies in Guinea pigs

Male Hartley GPs (200-250 gm/BW) at 8-10 weeks of age were obtained from National Laboratory Animal Center (NLAC), Taiwan, and maintained in the laboratory animal center of UBIA. All procedures on animals were performed in accordance with the regulations and guidelines approved by the Institutional Animal Care and Use Committee (IACUC) at UBIAsia. The GPs were vaccinated intramuscularly at weeks 0 and 3 with Montanide™ ISA50 V2-adjuvanted S1-RBD-based proteins. The animals received the primary dose of 200 μg of the vaccine (four injection sites, 0.25 mL/site) at week 0 and were boosted with 100 μg of the vaccine (two injection sites, 0.25 mL/site) at week 3. The immune sera from GPs (n = 5 for each protein immunogen) were collected at weeks 0, 3, and 5.

#### Adjuvant selection in rats

A total of 24 male Sprague Dawley rats at 8-10 weeks of age (300-350 gm/BW) were purchased from BioLASCO Taiwan Co., Ltd. After a 3-day acclimation, animals were randomly assigned to 8 groups. All procedures on animals were performed in accordance with the regulations and guidelines reviewed and approved by the IACUC at UBIAsia. The rats were vaccinated intramuscularly at weeks 0 (prime) and 2 (boost) with different doses ranging from 10 to 300 μg of UB-612 formulated in Montanide™ ISA 51 VG/CpG3 or Adju-Phos®/CpG1 adjuvant. The immune sera from rats (n = 3 for each dose group) were collected at weeks 0, 2, 3, and 4 for assessment of antigenic and functional activities. Local irritation analysis was conducted via a modified Draize technique [52] once daily within 24, 48 and 72 hrs after each vaccination. Additional abnormal clinical observations or lesions such as lameness, abscesses, necrosis, or local inflammation were recorded.

#### Rat Th1/Th2 balance study

A total of 12 male Sprague Dawley rats at 8-10 weeks of age (300-350 gm/BW) were purchased from BioLASCO Taiwan Co., Ltd. After a 3-day acclimation, animals were randomly assigned to 4 groups. All procedures on animals were performed in accordance with the regulations and guidelines reviewed and approved by the IACUC at UBIAsia. The rats were vaccinated intramuscularly at weeks 0 (prime) and 2 (boost) with three different doses (10 μg, 30 μg and 100 μg) of UB-612 formulated in Adju-Phos®/CpG1 adjuvant. The immune sera from rats (n = 3 for each dose group) were collected at weeks 0, 2, 3, and 4 for assessment of antigenic activities.

#### Vaccination and challenge procedure in AAV6/CB-hACE2 mice

A total of 12 male BALB/C at 8-10 weeks of age were purchased from BioLASCO Taiwan Co., Ltd. After a 3-day acclimation, animals were randomly assigned to 4 groups. All procedures on animals were performed in accordance with the regulations and guidelines reviewed and approved by the IACUC at UBI Asia. The mice were vaccinated by IM route at weeks 0 (prime) and 2 (boost) with 3, 9, or 30 μg of UB-612 formulated in Adju-Phos®/CpG1 adjuvant. The immune sera from mice were collected at weeks 0, 3 and 4 for assessment of immunogenic and functional activities by the assay methods described below.

AAV6/CB-hACE2 were produced by the AAV core facility in Academia Sinica (Taipei, Taiwan). BALB/C mice aged 8-10 weeks were anaesthetized by intraperitoneal injection of a mixture of Atropine (0.4 mg/ml)/Ketamine (20 mg/ml)/Xylazine (0.4%). The mice were then intratracheally (IT) injected with 3 x 10^11^ vg of AAV6/hACE2 in 100 μL saline. To transduce extrapulmonary organs, 1 x 10^12^ vg of AAV9/hACE2 in 100 μL saline was intraperitoneally injected into the mice.

Two weeks after AAV6/CB-hACE2 and AAV9/CB-hACE2 transduction, the mice were anesthetized and intranasally challenged with 10^6^ PFU TCID_50_ of SARS-CoV-2 virus (hCoV-19/Taiwan/4/2020 TCDC#4 obtained from National Taiwan University, Taipei, Taiwan) in a volume of 100 μL. Mice were weighed after the SARS-CoV-2 challenge daily. The mouse challenge experiments were evaluated and approved by the IACUC of Academia Sinica. Surviving mice were humanely euthanized in accordance with ISCIII IACUC guidelines.

#### Immunogenicity and protection studies in NHPs

Two non-human primate studies were conducted to evaluate the vaccination doses and numbers of immunizations, and the protective immunity against SARS-CoV-2 virus. All animal studies were approved by Institutional Animal Care and Use Committees (IACUC).

Animals were housed individually in stainless steel cages, an environmentally monitored, and well-ventilated room (conventional grade) maintained at a temperature of 18-26°C and a relative humidity of 40-70%. Animals were quarantined and acclimatized for at least 14 days. The general health of the animals was evaluated and recorded by a veterinarian within three days upon arrival. Detailed clinical observations, body weight, body temperature, electrocardiogram (ECG), hematology, coagulation and clinical chemistry were performed on the NHPs. The data were reviewed by a veterinarian before being transferred from the holding colony. Based on pre-experimental body weights obtained on Day -1, all animals were randomly assigned to respective dose groups using a computer-generated randomization procedure. All animals in Groups 1 to 4 were given either control or test article via intramuscular (IM) injection. Doses were administered to the quadriceps by injection of one hind limb. NHPs were observed at least twice daily (AM and PM) during the study periods for clinical signs which included, but were not limited to mortality, morbidity, feces, emesis, and changes in water and food intake. Animals were bled at regular intervals for the immunogenicity studies described below.

The first NHP study was conducted at JOINN Laboratories (Beijing) in Rhesus macaques aged approximately 3-6 years. Rhesus macaques were divided into four groups and injected intramuscularly with high dose (100 μg/dose), medium dose (30 μg/dose), or low dose (10 μg/dose) of vaccine or physiological saline. All grouped animals were immunized first at two time points (days 0, 28), and received a third boost dose at day 70. Blood samples were collected on days 0, 14, 28, 35, 42, 70 and 77 post-immunization. The macaques were challenged at 11 days after the 3^rd^ immunization (Day 81) with SARS-CoV-2 (10^6^ TCID50) intratracheally. The viral loads were determined by viral RNA copies/gram of lung tissue at 7 days after challenge (Day 88).

The second NHP study was conducted in cynomolgus macaques (3-6 years old). At Biomere animals were divided into three groups (5/group) and injected intramuscularly with high dose (100 μg/dose) or medium dose (30 μg/dose) of vaccine or physiological saline. All grouped animals were immunized at two time points (days 0, 28). At BIOQUAL blood samples were collected on days 0, 14, 28, 50 post-immunization. Four weeks after receipt of the final dose animals were challenged with a total of 1.0 × 10^5^ 50% tissue culture infectious dose (TCID_50_) of SARS-CoV-2 divided equally between intranasal (IN, 0.5 x 10^5^) and intratracheal (IT, 0.5 x 10^5^) administration. On days 0, 3, 5, and 8 post-challenge, bronchoalveolar lavage (BAL), nasal swabs and rectal swabs and were collected and viral load was assessed via RT-PCR.

### Immunoassay

#### ELISA for quantification of serum anti-S1-RBD antibody

Microtiter 96-well ELISA plates were coated with 2 µg/mL UBP Recombinant S1- RBD-His protein antigen (100 µL/well in coating buffer, 0.1 M sodium carbonate, pH 9.6) and incubated overnight (16 to 18 hrs) at 4°C. One hundred μL/well of serially diluted serum samples (10-fold diluted from 1:100 to 1:100,000, total of 4 dilutions) in 2 replicates were added and plates were incubated at 37°C for 1 hr. The plates were washed six times with 200 μL Wash Buffer (solution of phosphate buffered saline, pH 7.0-7.4 with 0.05% Tween 20 as surfactant). Bound antibodies were detected with standardized preparation of HRP-rProtein A/G (1:101 of Horseradish peroxidase-conjugated rProtein A/G dissolved in HRP-stabilizer) at 37°C for 30 min, followed by six washes with Wash Buffer. Finally, 100 μL of TMB (3,3’,5,5’-tetramethylbenzidine) prepared in Substrate Working Solution (citrate buffer containing hydrogen peroxide) was added into each well and incubated at 37°C for 15 min in the dark, and the reaction was stopped by adding 100 μL/well of Stop Solution (sulfuric acid solution, H_2_SO_4,_ 1.0 M). The absorbance at 450 nm was measured with an ELISA plate reader (Molecular Device, Model: SpectraMax M2e). The UBI® ELISA Titer Calculation Program was used to calculate the relative titer. The anti-S1-RBD antibody level was expressed as log_10_ of an end point dilution for a test sample (SoftMax Pro 6.5, Quadratic fitting curve, Cut-off value 0.5).

#### ELISA for binding inhibition of S1-RBD and human ACE2

96-well ELISA plates were coated with 2 µg/mL ACE2-ECD-Fc antigen (100 mL/well in coating buffer, 0.1M sodium carbonate, pH 9.6) and incubated overnight (16 to 18 hrs) at 4°C. The plates were washed 6x with Wash Buffer (25-fold solution of phosphate buffered saline, pH 7.0-7.4 with 0.05% Tween 20 (250 μL/well/wash) using an Automatic Microplate Washer. Extra binding sites were blocked by 200 μL/well of blocking solution (5 N HCl, Sucrose, Triton X-100, Casein, and Trizma Base). Two-fold serial dilutions (from 1:20 to 1:12,800) of immune serum or a positive control (diluted in a buffered salt solution containing carrier proteins and preservatives) were mixed with 1:100 dilution of S1-RBD- HRP conjugate (horseradish peroxidase-conjugated recombinant protein S1-RBD-His), incubated for 30±2 min at 25±2°C, washed and TMB substrate (3,3’,5,5’- tetramethylbenzidine diluted in citrate buffer containing hydrogen peroxide) was added. The reaction was stopped by adding stop solution (diluted sulfuric acid, H_2_SO_4,_ solution, 1.0 M) and the absorbance of each well was read at 450 nm within 10 min using the Microplate reader (VersaMax).

#### Flow cytometry assay for hACE binding inhibition

Human ACE2-transfected HEK293 cells (prepared in-house) were collected and washed with FACS buffer supplemented with 2% FBS (GIBCO, CN: 10099-148) in 1X PBS. One hundred μL of 20 μg/mL 2019-nCoV Spike Protein S1 (Fc Tag) (Sino Biological, CN: 40591-V02H) was mixed with 100 μL of antisera dilutions (serially 5-fold diluted from 1:5 to 1:3,125, total 5 dilutions) at 25°C for 1 hr. The mixture (200μL) was then added to the transfected cells (cell no. 2×10^5^) followed by incubation at room temperature for 1 hr. Cells were washed with FACS buffer and incubated with diluted (1:200) anti-human IgG Fc protein antibody (FITC) (Bethyl Laboratories, CN: A80-104F) on ice for an additional 30 min. After washing, the cells were analyzed in a FACSCanto II flow cytometry (BD Biosciences) using BD FACSDiva software.

#### Neutralization of live SARS-CoV2 by immune sera by CPE assay

Vero-E6 cells were expanded, and their concentrations were adjusted to 1.5x10^5^ viable cells/mL in culture medium (DMEM containing 10% FBS). The 96-well microtiter plates were seeded with 1.5x10^4^ cells/100 μL/well. The plates were incubated at 37°C in a CO_2_ incubator overnight. The next day, serum samples from vaccinated GPs were diluted (1:5) starting with 72 μL of serum sample + 288 μL of dilution medium (Dulbecco’s Modified Eagle Medium, DMEM, containing 5% FBS) to yield the first dilution. Then, 7x2- fold serial dilutions were made with dilution medium (dilution points were adjusted according to the characteristics of the sample). The challenge virus (SARS-CoV-2- TCDC#4, Taiwanese strain) was prepared at 100 TCID_50_ in 50 μL of culture medium, incubated with 50 μL volume of each serum dilution (50 μL) (in triplicates) for 1 hr at 37°C, before adding to Vero-E6 cells in triplicates. Medium only was co-incubated with an equal volume of 100 TCID_50_ of the viruses for 1 hr at 37°C and used as 100% infected control. The plates were incubated at 37°C in a CO_2_ incubator for 4 days. Cells were then fixed overnight with 100 μL of 10% formaldehyde prepared in phosphate buffered saline, pH 7.0-7.4, added into each well. The next day formaldehyde solution was discarded by inverting the plate, and 100 μL of crystal 0.5% violet staining solution was added into each well and incubated at room temperature for 1 hr. The infection rate was quantified by ELISA reader and image analysis. The infection rate of medium only at a challenge dose of 100 TCID_50_ virus was set at 100% and each serum dilution with greater than 50% infection was scored as infected. The 50% protective titer was determined by the Reed and Muench method [53, 54].

#### Neutralization assessment by immunofluorescence

Vero-E6 cell monolayers in 96-well plates from the CPE assay (described above) were also processed for visualization by immunofluorescence assay (IFA): the cells were stained with anti-SARS-CoV-2 N protein antibody and detected with anti-human IgG-488. The nuclei were counter stained with DAPI.

#### Neutralization assessment by plaque assay

Serum samples were heat-inactivated for 30 minutes at 56°C and diluted in a 2- fold serial fashion in MEM (Gibco) with Hepes (Corning) and Gentamicin sulfate (Cellgro) in U-bottom plates. 50 µL of each serum dilution was mixed with 50 PFU of virus in 50 µL. The serum/virus mixtures were incubated for 1 hr at 37°C. Fifty µl of the serum/virus mixtures were then transferred to Vero E6 cell monolayers in flat-bottom 96-well plates and incubated for 1 h at 37°C. The serum/virus mixture was then removed and replaced with 1:1 overlay composed of 1.1% methylcellulose (Fisher Chemical) and 2X MEM (Gibco) supplemented with gentamicin sulfate (Corning) and 4% FBS (Gibco). Plates were incubated 2 days at 37°C, then fixed with 10% neutral buffered formalin (Fisherbrand) according to approved SOP and removed from biocontainment. Viral plaques were counted after staining for 30 minutes with 1% crystal violet in formalin at room temperature.

#### ELISpot for measurement of cellular responses in rats

Spleens from vaccinated rats at 4 WPI were collected in Lymphocyte-conditioned medium (LCM; RPMI-1640 medium supplemented with 10% FBS and penicillin/streptomycin) and processed into single cell suspensions. Cell pellets were resuspended in 5 mL of RBC lysis buffer for 3 min at room temperature (RT), and RPMI- 1640 medium containing penicillin/streptomycin was then added to stop the reaction. After centrifugation, cell pellets were resuspended in LCM for use in the ELISpot assay. ELISpot assays were performed using the Rat IFN-γ ELISpot^PLUS^ kit (MABTECH, Cat. No.: 3220- 4APW), Rat IL-4 T cell ELISpot kit (U-CyTech, Cat. No.: CT081) and Rat IL-2 ELISpot Kit (R&D Systems, Cat. No.: XEL502). ELISpot plates precoated with capture antibody were blocked with LCM for at least 30 min at RT. 250,000 rat splenocytes were plated into each well and stimulated with S1-RBD-His protein plus Th/CTL peptide pool, S1-RBD-His protein, Th/CTL peptide pool, or each single Th/CTL peptide for 18-24 hrs at 37°C. Cells were stimulated with a final concentration of 1 μg of each protein/peptide per well in LCM. The spots were developed based on manufacturer’s instructions. LCM and ConA were used for negative and positive controls, respectively. Spots were scanned and quantified by AID iSpot reader. Spot-forming unit (SFU) per million cells was calculated by subtracting the negative control wells.

### Real-time RT-PCR for SARS-CoV-2 RNA quantification

The levels of N gene (subgenomic) mRNA (sgmRNA) were assessed by RT-PCR. The RT-PCR assay for the sgmRNA utilizes primers and a probe specifically designed to amplify and bind to a region of the N gene messenger RNA from SARS-CoV-2 (Primers: SG-N-F: CGATCTCTTGTAGATCTGTTCTC, SG-N-R: GGTGAACCAAGACGCAGTAT and Probe: FAM- TAACCAGAATGGAGAACGCAGTGGG -BHQ). A plasmid containing a portion of the N gene messenger RNA served as the control and semi-quantification standard. The cycling conditions were performed with a one-step PCR protocol: 48 °C for 30 minutes, 95 °C for 10 minutes followed by 40 cycles of 95 °C for 15 seconds, and 1 minute at 55 °C. Data were collected and calculated by Applied Biosystems 7500 Real-Time PCR System (Thermo Fisher Scientific, USA). The number of copies of RNA per mL of BAL or swab is calculated by extrapolation from the standard curve and multiplying by the reciprocal of 0.2 mL extraction volume to give a practical range of 50 to 5 x 10^7^ RNA copies per swab or mL BAL fluid.

To measure the RNA levels of SARS-CoV-2, specific primers targeting 26,141 to 26,253 regions in the envelope (E) gene of the SARS-CoV-2 genome were used by Taqman real-time RT-PCR previously described method [55]. Forward primer E-Sarbeco- F1 (5’-ACAGGTACGTTAATAGTTAATAGCGT-3’), reverse primer E-Sarbeco-R2 (5’-ATATTGCAGCAGTACGCACACA-3’), and probe E-Sarbeco-P1 (5’-FAM- ACACTAGCCATCCTTACTGCGCTTCG-BBQ-3’) were used. A total of 30 μL RNA solution was collected from each sample using a RNeasy Mini Kit (QIAGEN, Germany) according to the manufacturer’s instructions. Five microliters of RNA sample were added in a total mixture volume of 25 μL using Superscript III one-step RT-PCR system with Platinum Taq Polymerase (Thermo Fisher Scientific, USA). The final reaction mix contained 400 nM each of forward and reverse primers, 200 nM probe, 1.6 mM of deoxy- ribonucleoside triphosphate (dNTP), 4 mM magnesium sulphate, 50 nM ROX reference dye and 1 μL of enzyme mixture from the kit. The cycling conditions were performed with a one-step PCR protocol: 55°C for 10 min for cDNA synthesis, followed by 3 min at 94°C and 45 amplification cycles at 94°C for 15 sec and 58°C for 30 sec. Data were collected and calculated by Applied Biosystems 7500 Real-Time PCR System (Thermo Fisher Scientific, USA). A synthetic 113-bp oligonucleotide fragment was used as a qPCR standard to estimate copy numbers of viral genome. The oligonucleotides were synthesized by Genomics BioSci and Tech Co. Ltd. (Taipei, Taiwan).

### 50% tissue culture infectious dose (TCID_50_) assays

Mouse lung tissues were weighed and homogenized in 1 mL of DMEM with 1% penicillin/streptomycin using a homogenizer. After centrifugation at 13,000 rpm for 10 min, supernatant was harvested for live virus titration (TCID_50_ assay). Briefly, serial 10-fold dilutions of each sample were inoculated in a Vero-E6 cell monolayer in quadruplicate and cultured in DMEM with 1% FBS and penicillin/streptomycin. The plates were observed for cytopathic effects for 4 days. TCID_50_ was interpreted as the amount of virus that caused cytopathic effects in 50% of inoculated wells. Virus titers were expressed as TCID50/mL.

### Histopathological analysis of lung tissues

After fixation with 10% formaldehyde for one week, lung tissues were trimmed, processed, embedded, sectioned and stained with Hematoxylin and Eosin (H&E), followed by microscopic examination. To score the lung histopathology, lung section was divided into 9 equal square areas using a 3 × 3 grid. Lung tissue of every area was scored using a scoring system. Scores from each of the 9 areas were averaged and this average value was designated as the animal’s score.

The scoring system was as follows: 0, Normal, no significant finding; 1, Minor inflammation with slight thickening of alveolar septa and sparse monocyte infiltration; 2, Apparent inflammation, alveolus septa thickening with more interstitial mononuclear inflammatory infiltration; 3, Diffuse alveolar damage (DAD), with alveolus septa thickening, and increased infiltration of inflammatory cells; 4, DAD, with extensive exudation and septa thickening, shrinking of alveoli, restricted fusion of the thick septa, obvious septa hemorrhage and more cell infiltration in alveolar cavities; 5, DAD, with massive cell filtration in alveolar cavities and alveoli shrinking, sheets of septa fusion, and hyaline membranes lining the alveolar walls.

## Acknowledgements

We thank Dr. Qian Gao of Sinovac for providing CPE neutralization titrations free of charge.

