## Supplemental Information for "A Novel SARS-CoV-2 Multitope Protein/Peptide Vaccine Candidate is Highly Immunogenic and Prevents Lung Infection in an AAV hACE2 Mouse Model and non-human primates"

A

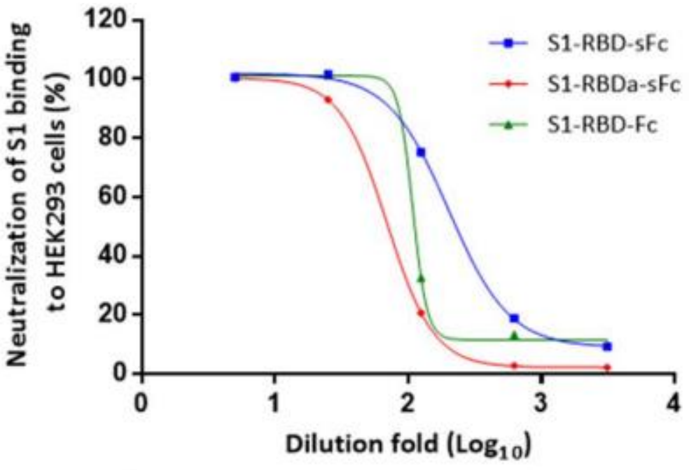


B

Dilution $ID_{50}$ values

| Immunogens | $ID_{50}$ |
|---|---|
| S1-RBD-sFc | 202 |
| S1-RBDa-sFC | 69.2 |
| S1-RBD-Fc | 108 |

**Figure S1.** Neutralization and dilution ID50 in S1 protein binding to ACE2-expressing HEK293 cells by guinea pig immune sera pooled at 2 weeks after the 2nd immunization (Week 5). Serum samples from each vaccinated animal were pooled and serially diluted for assay of inhibition of S1 protein binding to the ACE2-expressing HEK293 cells. The inhibition of binding to the cells was analyzed on flow cytometry (a) and the 50% neutralization titer (ID50) (b) determined based on the inhibition curve generated by four-parameter logistic regression.

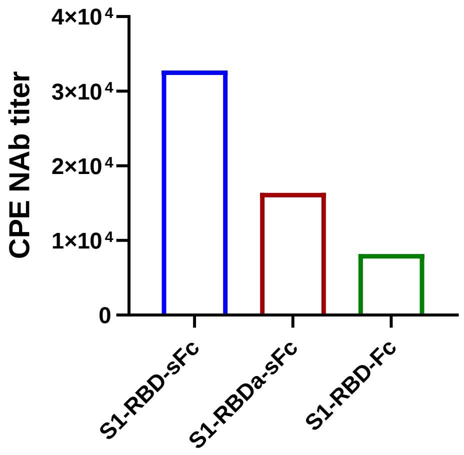


**Figure S2.** Neutralizing antibodies determined by CPE assay in group-pooled guinea pig sera collected at 2 weeks after the 2$^{nd}$ immunization (Week 5).

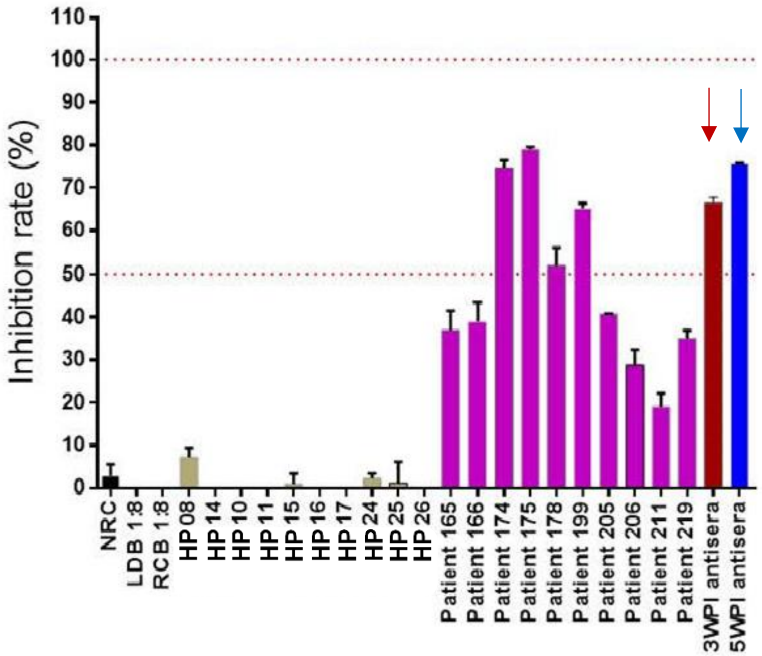


**Figure S3. Comparative S1-RBD:hACE2 binding inhibition by GP sera and convalescent sera.** SARS-CoV-2 inhibition rates were evaluated with human serum samples from normal healthy persons (HP, n=10) and virologically diagnosed COVID-19 patients (n=10) tested at a 1:20 dilution. Pooled immune sera from S1-RBD-sFc vaccinated GP collected at Week 3 (3WPI) and Week 5 (5WPI) indicated with arrows, were tested at 1:1000 and 1:8000 dilutions, respectively. NRC was served as non-reactive control.

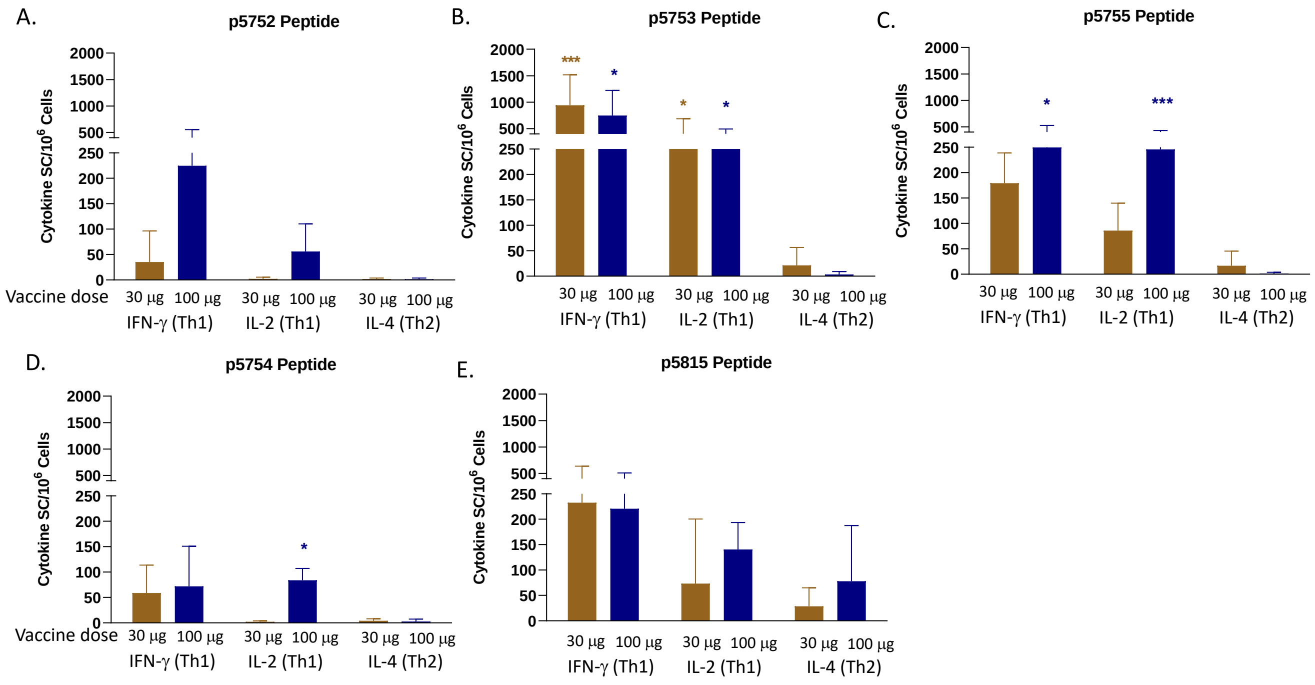


**Figure S4. Cellular immunogenicity testing in rats (ELISpot detection of IFN-γ, IL-2 and IL-4 secreting cells in UB-612 immunized rats).** Groups of rats were immunized with 30 µg or 100 µg UB-612 vaccine at Weeks 0 and 2. Splenocytes were collected at Week 4 (2 weeks after the 2nd immunization) and stimulated with individual peptides, panels A-C: S2 peptides (p5752, p5753 and p5755); panel D: N peptide (p5754); panel E: M peptide (p5815). IFN-γ, IL-2 and IL-4-secreting splenocytes were determined by ELISpot analysis. Cytokine-secreting cells (SC) per million cells was calculated by subtracting the negative control wells. Bars represent the mean SD (n = 3). The secretion of IFN-γ or IL-2 was observed to be significantly higher than that of IL-4 in the 30 and 100 µg group (* $p < 0.05$, ** $p < 0.01$, *** $p < 0.005$).

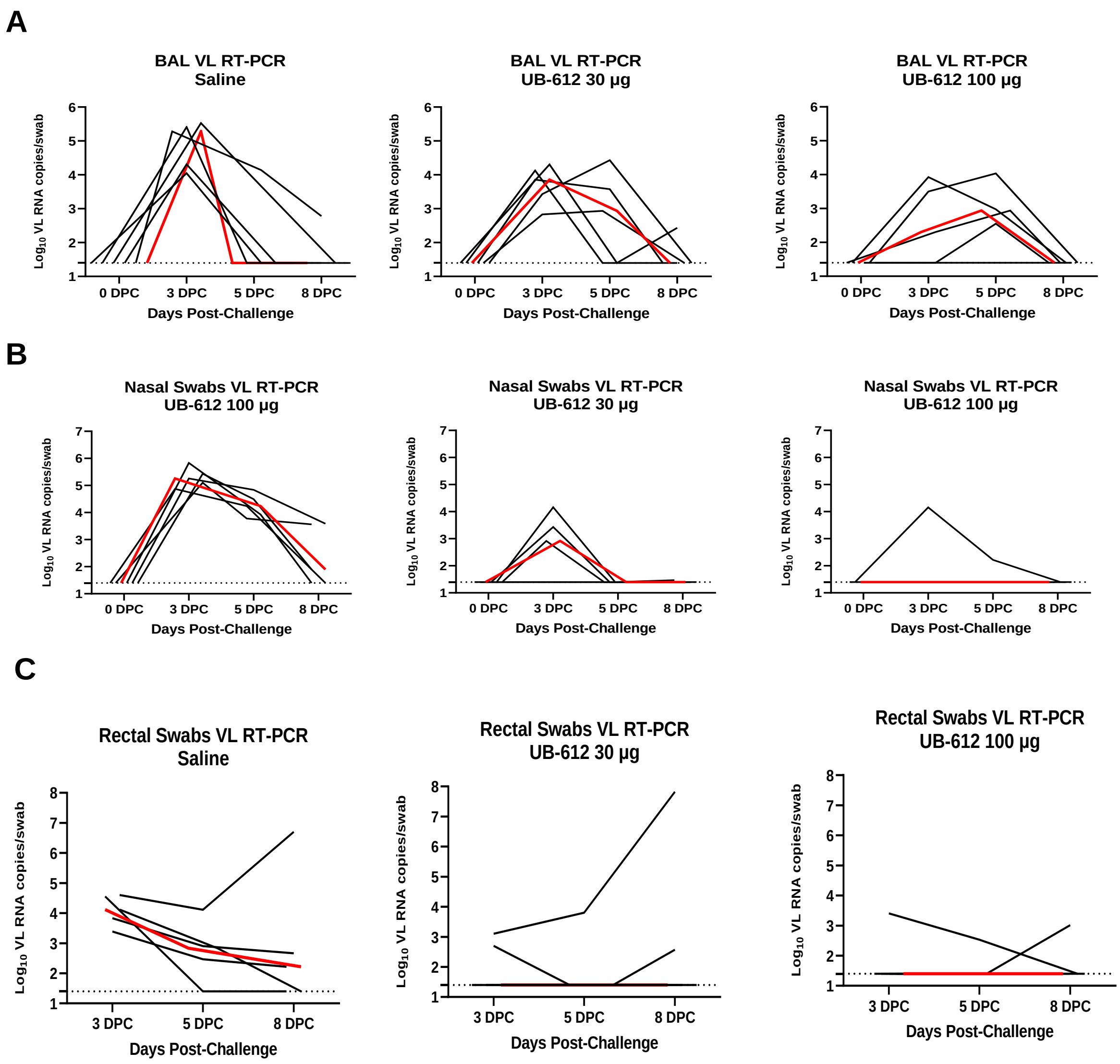


**Figure S5. Protective immunity in rhesus macaque Study 2.** The 3 groups of macaques (5/group) received Saline, UB-612 30 μg or UB-612 100 μg, respectively, on Day 0 and Day 28, and challenged with SARS-CoV-2 virus on Day 55. The viral loads in each macaque were detected in BAL (A), nasal swaps (B) and rectal swaps (C) by VL RT-PCR at 3, 5 and 8 days after challenge. The black curves represent viral loads of the individual animals and the red curves are the median viral load of each group.
